# Contextualising the developability risk of antibodies with lambda light chains using enhanced therapeutic antibody profiling

**DOI:** 10.1101/2023.06.28.546839

**Authors:** Matthew I. J. Raybould, Oliver M. Turnbull, Annabel Suter, Bora Guloglu, Charlotte M. Deane

## Abstract

Antibodies with lambda light chains (*λ*-antibodies) are generally considered to be less developable than those with kappa light chains (*κ*-antibodies), leading to substantial systematic biases in drug discovery pipelines. This has contributed to kappa dominance amongst clinical-stage therapeutics. However, the identification of increasing numbers of epitopes preferentially engaged by *λ*-antibodies shows there is a functional cost to neglecting them as potential lead candidates during discovery campaigns. Here, we update our Therapeutic Antibody Profiler (TAP) tool to use the latest data and machine learning-based structure prediction methods, and apply this new protocol to evaluate developability risk profiles for *κ*-antibodies and *λ*-antibodies based on their surface physicochemical properties. We find that *λ*-antibodies are on average at a higher risk of poor developability — as an indication, over 40% of single-cell sequenced human *λ*-antibodies are flagged by TAP for risk-prone patches of surface hydrophobicity (PSH), compared to around 11% of human *κ*-antibodies. Nonetheless, a substantial proportion of natural *λ*-antibodies are assigned more moderate risk profiles by TAP and should therefore represent more tractable candidates for therapeutic development. We also analyse the populations of high and low risk antibodies, highlighting opportunities for strategic design that TAP suggests would enrich for more developable *λ*-based candidates. Overall, we provide context to the differing developability of *κ*- and *λ*-antibodies, enabling a rational approach to incorporate more diversity into the initial pool of immunotherapeutic candidates.

## Introduction

Antibodies are the dominant category of biotherapeutics; more than 140 therapeutic antibodies have now been approved by regulators with over 550 currently active in clinical trials (1, 2). Their popularity is tied to their use by natural immune systems and their ability to achieve high affinity/specificity for seemingly any targeted pathogen (‘antigen’), enabling its selective eradication (3).

Antibodies accomplish precise antigen recognition through two dedicated antigen binding sites, termed the variable regions (Fvs). These Fvs are identical and structurally/chemically intricate, comprising six proximal complementarity-determining region (CDR) loops spread across two polypeptide chains — three on the variable domain of the heavy chain (VH, CDRH1-3) and three on the variable domain of the light chain (VL, CDRL1-3).

VH is encoded across three genes (a combination of a heavy V, D, and J gene) while VL is encoded across two genes (a combination of a light V and J gene). Within each locus (chromosome region that contains a certain category of immunoglobulin gene), there exist many alternative chemically-distinct ‘germline’ genes (4). During development, V(D)J gene recombination events occur largely at random, creating considerable combinatorial diversity (5). Nucleotide insertion/deletions in the junction region between genes (which falls within the CDR3 loops) further contribute to exceptional sequence diversification, magnified naturally through somatic hypermutation during an immune response or artificially through *in vitro* affinity maturation/engineering.

While the heavy chain V, D, and J genes (IGHV, IGHD, IGHJ) lie solely on chromosome 14, light chain V and J genes exist at two genetic loci; a ‘kappa’ (*κ*, IGKV and IGKJ) locus on chromosome 2, and a ‘lambda’ (*λ*, IGLV and IGLJ) locus on chromosome 22.

Light chain identity is often critical to an antibody’s function. For example, it has been observed in different toxin, virus, and vaccine response contexts that *κ*- and *λ*-antibodies are expressed in characteristic proportions with restricted usages, and that they tend to have different antigen specificities (6). Amongst the thousands of anti-coronavirus antibodies independently isolated throughout the pandemic, the same light chain gene origins have been frequently observed amongst antibodies with a high confidence of engaging the same epitope (7, 8). This link between light chain gene and function has recently been shown to apply more generally, as evidenced by Jaffe *et al*. who found ‘light chain coherence’ of memory B-cell compartments (9), and by Shrock *et al*. who identified the presence of germline amino acid-binding motifs — many of which lie on the light chain (10). Together, these phenomena are likely by-products of the documented sequence (10–13) and structural (10, 14–16) differences between *κ*- and *λ*-VLs, which may have evolved to increase the efficacy of receptor editing (17), a process during which maturing BCRs can exchange their initial recombined *κ*-VLs for *λ*-VLs to prevent autoreactivity.

Despite their functional utility, *λ*-antibodies are currently under-represented across clinical-stage therapeutic antibodies (CSTs). Of a set of 242 CSTs curated in 2019 (18), all of which were designed for human application, only around 10% derived from *λ*-genes. By comparison, *λ*-antibodies are estimated to comprise roughly 35% of natural human repertoires (19, 20).

The precise reasons behind the paucity of *λ*-antibodies in the clinic are unknown, but there are several probable origins. These include factors related to the dominant methods of therapeutic discovery (21, 22), such as unintended selection bias in screening library design (23) and the higher *κ*:*λ* ratios (up to 20:1) seen in mouse antibody repertoires (24).

There is also evidence that *λ*-antibodies fail more frequently than *κ*-antibodies to advance through pre-clinical development (25). Several studies have identified that *λ*-VLs exhibit a higher average hydrophobicity than *κ*-VLs (11, 12, 17, 18); higher hydrophobicity suggests an increased propensity towards the formation of aggregates *via* the hydrophobic effect. This mechanism is understood to be the primary force driving light chain amyloidosis, where free light chains self-associate, and data suggests that *λ*-VLs prone to dimerisation outnumber *κ*-VLs (26).

This has earned *λ*-antibodies a reputation for poor developability that has fed back into systematic discovery biases, such as through the intentional development of *κ*-only screening libraries (27), or, when given a choice of progressing *κ*- or *λ*-antibodies to downstream lead optimisation, a tendency to prioritise the former. However, it is probable that a sizeable proportion of *λ*-antibodies are indeed developable, and that rational engineering could be used to make some more challenging *λ*-antibodies biophysically tractable (28). In general, better distinction between more developable and less developable *λ*-antibodies should be applied to limit the degree to which we artificially restrict candidate diversity, and therefore targetable epitope space, during early stage discovery.

In 2019, we published the Therapeutic Antibody Profiler (TAP), a method for the computational developability assessment of lead candidates based on comparing their 3D biophysical properties to those of CSTs (18). At the time, we only had access to 25 *λ*-based CST sequences and artificially-paired representations of natural human antibodies. Now, through dedicated efforts to track the sequences of CSTs as they are designated by the World Health Organisation (WHO) (1) and increased public availability of paired-chain natural antibody repertoires (29), we are able to more confidently profile the physicochemical properties of therapeutic and natively-expressed human *λ*-antibodies.

In this paper, we first improve TAP by incorporating ABodyBuilder2 (30), a state-of-the-art deep-learning based antibody structure prediction method and highlight changes and robustness of the new guideline values. We then use our updated protocol to characterise developability-linked biophysical differences across CST and natural *κ*-and *λ*-antibodies. Finally, we probe the subset of red-flagging antibodies for recurrent features associated with extreme scores, and which may be avoided to derisk the incorporation of *λ*-antibodies into screening libraries.

Overall, our study provides an improved methodology for therapeutic antibody profiling and adds context to the developability of *λ*-antibodies, facilitating their selective consideration as leads during early-stage drug discovery.

## Results

### Curating Datasets of Therapeutic and Natural Antibodies

We first curated the latest set of non-redundant, post Phase-I clinical stage therapeutics (CSTs) designated for use in humans from Thera-SAbDab (1) (25^th^ January, 2023). We obtained 664 CST Fv sequences (the **‘CST**_**all**_**’** dataset), compared to the 242 used in our previous analysis (18). We also obtained all 79,759 non-redundant natively-paired human antibody sequences from the new Observed Antibody Space database (29) (25^th^ January, 2023); the **‘Nat**_**all**_**’** dataset. This compares to datasets of between 14,000-19,000 artificially-paired human antibody sequences used in our previous analysis (18).

### Benchmarking a New TAP Modeling Protocol

The original Therapeutic Antibody Profiler used the homology modeling tool ABodyBuilder (31) (‘ABodyBuilder1’, for clarity) for antibody structural modeling. In 2018, this was the state-of-the-art tool for high-throughput antibody modeling. However, recent advances in deep learning have yielded several pretrained *ab initio* structure prediction architectures that can be applied or adapted to the task of rapid antibody/CDR loop modeling (30, 32–34). Their average performance has been shown to be consistently higher than that of homology-based antibody modeling methods. Since better models of antibodies should improve the reliability of our developability guidelines, we explored the case for updating our TAP protocol to use a more recent machine learning-based tool (ABody-Builder2 (30)) for 3D structural modeling.

We first confirmed that ABodyBuilder2’s improved general performance translates specifically to CSTs, observing increased backbone and side chain modeling accuracy relative to ABodyBuilder1 across a set of recently-solved therapeutics (see Supporting Information Results, Supporting Information Methods, Tables S1-S2). This motivated us to formally adopt ABodyBuilder2 as the tool for 3D modeling prior to computation of the TAP metrics.

We next analysed the impact of using ABodyBuilder2 *versus* ABodyBuiler1 for structural modeling on the TAP developability guidelines calculated across the CST_all_ set. We measure this based on their impact on the amber and red flagging thresholds — characteristic percentile values used to demark the extrema of each TAP property distribution linked to poor developability (18).

For reference, amber flags for Total CDR Length (L_tot_) or Patches of Surface Hydrophobicity (PSH) are assigned to scores in the 0^th^-5^th^ or 95^th^-100^th^ percentiles relative to CSTs, while red flags are assigned if the L_tot_/PSH score falls below the 0^th^ or above the 100^th^ percentile. Amber flags for Patches of Positive Charge (PPC) or Patches of Negative Charge (PNC) are assigned if a score falls in the 95^th^-100^th^ percentile range relative to CSTs, and red flags are assigned to scores above the 100^th^ percentile. Finally, amber flags are assigned for the Structural Fv Charge Symmetry Parameter (SFvCSP) metric if the score falls between the 0^th^-5^th^ percentile values relative to CSTs, while red flags are assigned to scores below the 0^th^ percentile.

The ABodyBuilder2-modeled CST_all_ flagging thresholds show high similarity to those obtained by ABodyBuilder1 (Table 1). There is some evidence of a systematic bias associated with the different modeling protocol. Comparing the amber flag thresholds (less volatile than red flag thresholds as they capture 5% of the data) shows that ABodyBuilder2-modeled CSTs have lower PSH scores than ABodyBuilder1-modeled CSTs. This drop in PSH score is consistent with ABodyBuilder2’s more accurate modeling; we found in our original TAP paper that PSH values calculated over solved crystal structures (theoretically ‘perfect’ predictions) were lower on average (a difference of *c*. 10) than those calculated over ABodyBuilder1 models of the same CSTs (18).

**Table 1.** Flagging regions across the five TAP developability metrics calculated over the CST_all_ dataset (See Methods), when therapeutics are modeled by ABodyBuilder1 or by ABodyBuilder2. As ABodyBuilder1 failed to produce a model for Basiliximab, Iscalimab, and Netakimab, its statistics are calculated over 661/664 CSTs. Amber and red flag percentiles were set as per Raybould *et al*. 2019 (18). L_tot_: Total CDR Length, PSH: Patches of Surface Hydrophobicity metric, PPC: Patches of Positive Charge metric, PNC: Patches of Negative Charge metric, SFvCSP: Structural Fv Charge Symmetry Parameter.

| TAP Property | ABodyBuilder1 |  | ABodyBuilder2 |  |
| --- | --- | --- | --- | --- |
|  | Amber Flag Region | Red Flag Region | Amber Flag Region | Red Flag Region |
| $L_{tot}$ | 37 to 42<br>55 to 63 | < 37<br>> 63 | 37 to 42<br>55 to 63 | < 37<br>> 63 |
| PSH | 90.88 to 114.61<br>174.26 to 219.20 | < 90.88<br>> 219.20 | 95.58 to 110.11<br>168.06 to 201.59 | < 95.58<br>> 201.59 |
| PPC | 1.27 to 4.21 | > 4.21 | 1.32 to 4.22 | > 4.22 |
| PNC | 1.93 to 4.36 | > 4.36 | 2.00 to 4.42 | > 4.42 |
| SFvCSP | -30.00 to -6.00 | < -30.00 | -30.60 to -6.00 | < -30.60 |

### Testing the Robustness of the TAP Developability Guidelines

We then probed the robustness of our developability guidelines to various perturbations. TAP values calculated on the subset of CSTs modeled with higher certainty should be more reliable. Model confidence can be estimated through the frame-aligned prediction error (FAPE) metric, a property minimised as part of the ABodyBuilder2 loss function that can be interpreted as a measure of backbone prediction uncertainty for each residue (30). To inves- tigate the impact of FAPE-based confidence filtering on our guidelines, we first determined an appropriate CDRH3 root-mean squared predicted error threshold that would filter out the least-confidently modeled CDRH3s (1.31 Å, see Methods for the derivation), then calculated our developability guidelines based only on the subset of most confident CST predictions (the **CST**_**conf**_ set, see Fig. S1, Table S3). Overall, this filtering had only a small impact on the guidelines, with a slight downsampling of the numbers of CSTs with longer CDRH3s and higher PSH, suggesting that the new TAP guidelines are robust to the variable prediction accuracy within ABodyBuilder2.

Next, we examined the effect of ABodyBuilder2’s non-deterministic side chain modeling to explore statistically how side chain conformational uncertainty impacts the guidelines.

We ran the TAP protocol three times per CST and investigated the consistency of structure-dependent TAP metrics for each CST (Fig. S2). The results for all metrics were all highly consistent between runs. PPC, PNC, and SFvCSP values were the most consistent, with Pearson’s coefficient values in the range of 0.993-0.996. Due to their sensitivity to structural variations in any CDR vicinity residue, we expected the PSH values to be more susceptible to inter-run fluctuations. This was borne out, however PSH remained strongly correlated between two independent modeling runs (*ρ*: 0.945), and even more strongly correlated between one run and the mean of three independent runs (*ρ*: 0.981). The proportions of flagging inconsistencies (instances where a CST would be flagged for that property based on one TAP run but not based on three repeats), were as follows: PSH (lower): 3.31%, PSH (upper): 1.81%, PSH (overall): 5.12%, PPC: 0.30%, PNC: 0.30%, SFvCSP: 0.75%.

To capture the absolute variability of scores across repeats, we evaluated for each metric/CST the variance across the three runs and averaged these values on a per metric basis across the CST_all_ dataset. The mean PSH variance was 10.53 while the mean PPC, PNC, and SFvCSP variances were below 1 (Table S4). When values from three runs were amalgamated to establish aggregate TAP guidelines, this translated to a very small variation in threshold values from those obtained based on a single model of each CST (Table S5).

In addition to this statistical sampling of energy-minimised side chain conformations, we also evaluated the variation in TAP scores calculated over the course of molecular dynamics simulations; incorporation of dynamics in guideline evaluation was suggested in a recent study on computational developability prediction (35). We selected 14 case study CSTs, seven of which had solved Fv coordinates in the ABody-Builder2 training set and seven of which did not; for details of the molecular dynamics simulation and TAP calculations, see Methods.

The profiles for each of the four structure-dependent TAP metrics are shown in the Supporting Information (Figs. S3-S6). The mean value of the TAP properties over the course of the simulation showed good agreement with an ensemble of three TAP predictions on the static Fv models output directly by ABodyBuilder2. Based on a paradigm where if a developability flag is raised on any of the repeat calculations we consider the antibody formally flagged for that property, the ensemble of direct ABodyBuilder2 outputs agreed with the flag assigned to the simulation mean for 12/14 calculations for PSH, 13/14 calculations for PPC, and 14/14 calculations for PNC and SFvCSP.

### TAP metric profiles over time and by development stage

Finally, we investigated the impact on our metric distributions of filtering our CSTs by metadata properties.

To assess the properties of CSTs over time, we split the set by the year they were given a proposed WHO International Non-proprietary Name, yielding 356 named between 1987-2017 and 308 named between 2018 and the present day. Comparing their TAP property distributions (Fig. S7) indicates that while their charge metrics are similar, the amber and red flag thresholds of the L_tot_ and PSH properties have shifted to more extreme values at both tails, suggesting an increased willingness to push CST design into new property spaces and perhaps reflecting formulation advances able to accommodate more extreme physicochemical properties.

Recent studies have suggested that developability guidelines may be better derived from marketed therapeutics (35, 36); we evaluated our TAP distributions for the subsets of CSTs in Phase-II (341), Phase-III (141), or in Preregistration/Approved (182), however observed no clear trend in their properties along the clinical progression axis (Fig. S8). Equally, we saw little difference in the properties of CSTs known to be in active development or that completed the development pipeline *versus* CSTs whose campaigns were terminated before reaching approval (Fig. S9). These observations are consistent with the principle that CSTs with unmanageable developability issues do not tend to progress past pre-clinical/Phase-I development, and that decisions to terminate campaigns at later clinical stages are often attributable to other causes.

### Updated comparison of CSTs to natural human antibodies

A key biotechnological advance in recent years has been the advent of high-throughput paired B-cell receptor (BCR) sequencing (37). Publicly available paired antibody sequences are increasingly abundant (29) and provide a higher fidelity comparison set than the artificially-paired natural single chain reads we used in previous repertoire characterisation work (18, 38). These samples, coupled with the availability of 2.75 times as many CSTs and a more accurate/versatile modeling protocol, provides an ideal opportunity to revisit prior analyses and explore whether we observe similar trends in the biophysical properties of therapeutics and natural antibodies.

We calculated the TAP profiles for our new curated datasets of naturally-paired human sequences (Nat_all_) and CSTs (CST_all_). The patterns of the distributions aligned with our findings in the original paper (Fig. 1A-E). CSTs and natural human antibodies adopted similar PPC, PNC, and SFvCSP distributions, but natural antibodies were even more enriched at longer CDRs and higher PSH scores than observed previously (30.16% and 24.23% fall above the upper amber flag thresholds set by the top-5% of CSTs, respectively). To ensure the length bias was not the sole driver of higher PSH scores, we plotted the L_tot_ against the PSH score for the Nat_all_ and CST_all_ datasets (Fig. 1F). While almost all the natural antibodies found at extreme L_tot_ values flag for PSH, so too do a disproportionate number of natural antibodies at more moderate CDR lengths, even down to the smallest recorded L_tot_ value.

**Fig. 1.**
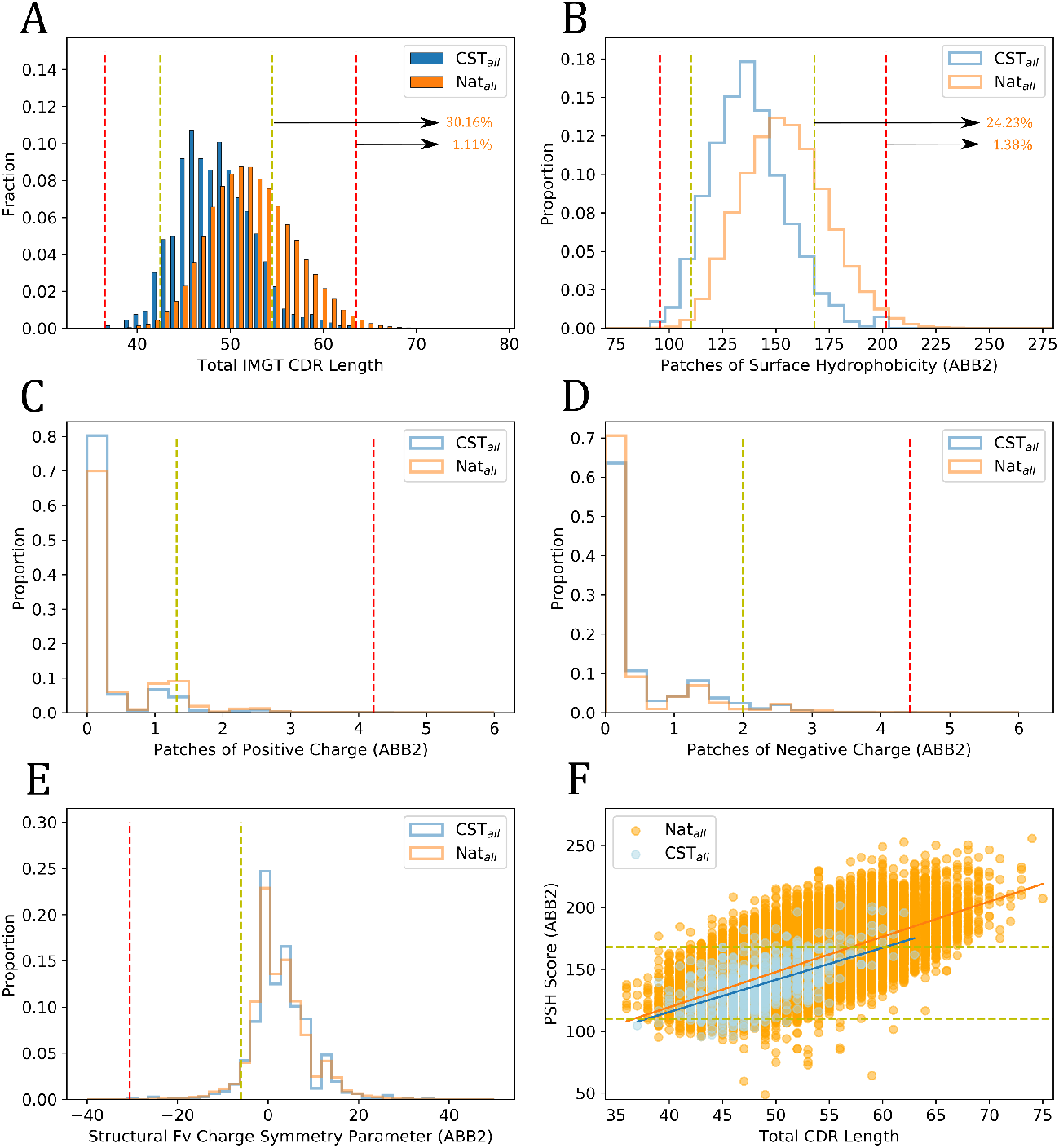
The five TAP developability metric distributions set by using ABodyBuilder2 (30) (ABB2) to model the curated CST_all_ (blue) and Nat_all_ (orange) datasets. An additional plot displays the trend between L_tot_ and PSH Score for both datasets. Amber and red flag thresholds are shown with dashed lines of corresponding color. The percentages of Nat_all_ antibodies lying above the L_tot_ and PSH upper thresholds are highlighted.

To further test the robustness of these conclusions, we then restricted our analysis to a confidence-filtered subset of ABodyBuilder2 models of the CSTs (CST_conf_) and the natural data (**Nat**_**conf**_), generated using the FAPE threshold bench-marked on CSTs (see Methods). In these sets a much smaller fraction of natural antibodies survived the filtering cut-off (∼38% of Nat_all_ *versus* ∼75% of CST_all_), likely due to the fact that natural antibodies sample longer CDRH3 lengths — which are both more conformationally diverse and harder to crystallise — as well as the general under-representation of natural antibodies in the Protein Data Bank (39), on which ABodyBuilder2 is trained.

The resulting CST_conf_ and Nat_conf_ TAP distributions show analogous relative positioning to our original results (18), with CSTs occupying shorter L_tot_ and smaller PSH values than natural antibodies, but having similar charge characteristics (Fig. S1). Quantitatively, over 16% and over 18% of Nat_conf_ antibodies surpassed the L_tot_ and PSH upper amber thresholds set by the CST_conf_ set (compared with *∼*30% and *∼*24% on the Nat_all_ set, respectively). The large reduction in the number of natural antibodies flagging for L_tot_ confirms that antibodies with longer CDR loops are modeled with lower confidence. The smaller percentage reduction in natural antibodies flagging for PSH reflects the increased tendency for natural antibodies of all lengths to occupy higher PSH values.

In summary, our investigations strengthen the evidence that CSTs and natural antibodies differ in key physicochemical properties.

### Using the new TAP protocol to explore *λ*-antibody developability

We then used our updated TAP protocol to explore the relative developability of *κ*- and *λ*-antibodies.

We examined the growth trends of *κ*- and *λ*-CSTs. Plotting their numbers over time reveals distinct patterns in usage (Fig. 2). For example, while novel *κ*-CST Fvs have been continuously released in double-digit quantities per year since 2007, new *λ*-CST Fvs only reached this level in 2018. In 2019, for example, the industry developed 53 new *κ*-CST Fvs, but only 10 new *λ*-CSTs.

**Fig. 2.**
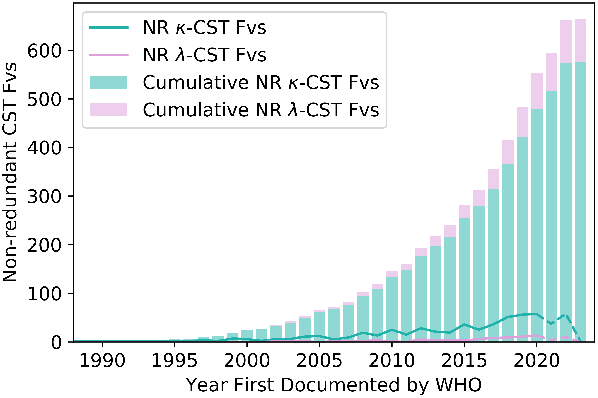
Tracking the numbers of 100% sequence non-redundant *κ* and *λ* variable regions (Fvs) across post Phase-I sequence non-redundant clinical stage therapeutics (NR CSTs) from 1988 to 2023. The x-axis reflects the year in which each CST was granted a proposed International Non-proprietary Name (INN) by the World Health Organisation (WHO). Cumulative totals are shown through a stacked bar chart, while year-by-year totals are shown in the line graph. 2021-2023 are shown in dashed lines; these totals will likely change significantly once more therapeutics first reported in these years have had time to advance past Phase-I Clinical Trials.

This lag has led to a significant disparity in the abundance of *κ*- and *λ*-CSTs. As of January 2023, Thera-SAbDab contained 576 non-redundant *κ*-CST Fvs (86.7%) and 88 non-redundant *λ*-CST Fvs (13.3%); far below the relative abundance of human *λ*-antibodies (30-35% (19, 20)). However, prior to the disruption of the pandemic, there were signs of an upwards growth trend in *λ*-CSTs (Fig. 2). There is evidence to suggest this is driven by the propensity *λ*-VLs to bind different targets/epitopes to *κ*-VLs; amongst the therapeutics designated by the WHO since 2022, six *λ*-antibodies (Acimtamig, Firastotug, Golocdacimig, Temtokibart, Tolevibart, and Zinlivimab) are first-in-class clinical candidates against novel antigen targets or epitopes (FCGR3A, HHV gB AD, OLR1, IL22RA1, HPV Envelope Protein, and the HIV-1 gp120 V3 epitope, respectively (1)). Overall, the 88 sequence non-redundant *λ*-CSTs now in Thera-SAbDab represents a 250% increase on the 25 *λ*-CSTs we had access to when developing our original guidelines.

### TAP distributions across *κ*- and *λ*-antibodies

The CST biophysical property distributions for the two types of light chain are shown in Fig. 3A-E. *λ*-CSTs disproportionately amber flag at the upper extrema of the L_tot_ (3.1% *κ*, 27.27% *λ*) and PSH (2.7% *κ*, 21.1% *λ*) distributions, and, to a lesser extent, for PPC (4.1% *κ*, 11.8% *λ*). In contrast, *κ*-CSTs predominate in the lower extrema of the L_tot_ (10.4% *κ*, 1.3% *λ*) and PSH (5.8% *κ*, 0% *λ*) distributions. While the relative proportions in the flagging region for the SFvCSP score are similar, only *κ*-CSTs occupy the most extreme values (below -15).

**Fig. 3.**
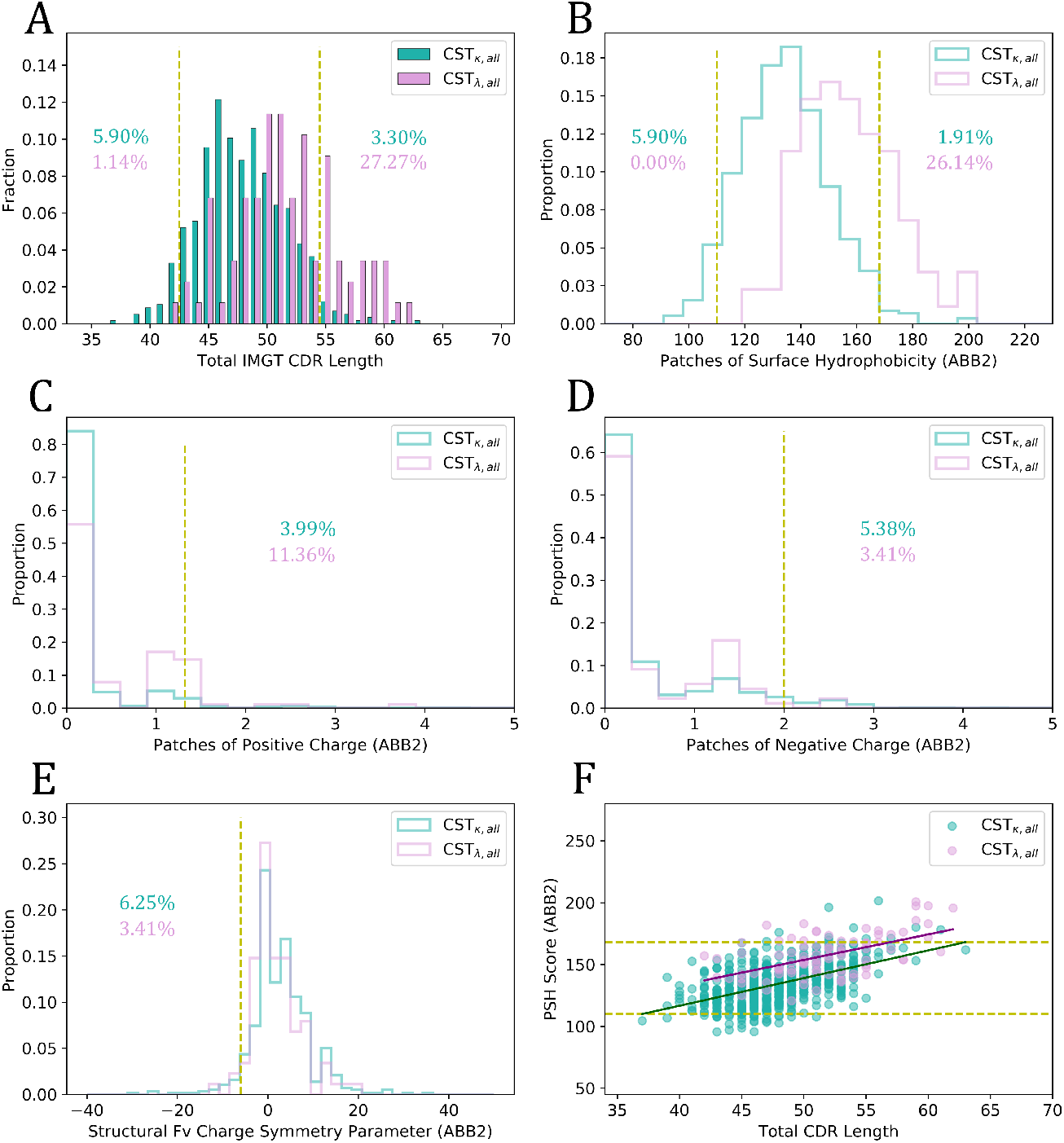
Plots of the five TAP properties for the *κ* (seagreen) and *λ* (plum) CSTs, and the trend of the Patches of Surface Hydrophobicity (PSH) score with L_tot_, using ABodyBuilder2 for structural modeling (ABB2). Amber thresholds are set based on the 5^th^ and/or 95^th^ percentile values of the combined set of kappa and lambda CSTs. Percentage values reflect the proportions of the correspondingly coloured light-chain class of antibody within the amber-flagged region of each distribution.

As PSH values correlate to some extent with L_tot_, we checked whether the preponderance of *λ*-CSTs at high PSH values was simply a by-product of length (Fig. 3F). Our results indicate that *λ*-CSTs are not significantly more driven towards high PSH scores by longer CDR lengths than *κ*-CSTs are, with *λ*-CSTs having higher average PSH scores than *κ*-CSTs at every sampled L_tot_ value.

The observation that *λ*-CSTs sit at such longer average L_tot_ values than *κ*-CSTs was surprising. While *λ*-CDRL3s are known to be longer on average than their *κ* equivalents (17), this alone cannot explain the shift. Instead, for this dataset, the disparity is also driven by biased pairing of *λ*-VLs with VH chains with longer average CDRH3 lengths (*μ*_*κ*-CST, H3_: 12.53 *±* 3.07), *μ*_*λ*-CST, H3_: 14.30 *±* 3.87).

We then studied the biophysical property distributions for the natural human sets of *κ*- and *λ*-antibodies (Fig. 4). On these datasets we found a much smaller difference in L_tot_ scores between the natural *κ*-antibody and *λ*-antibody models than observed in the CSTs; an offset fully explained by CDRL3 length biases across the two types of light chain (*μ*_*κ*-Nat, L3_: 9.12 ± 0.8, *μ*_*λ*-Nat, L3_: 10.61 ± 1.03). This result is consistent with Townsend *et al*. (17) and provides strong evidence that the longer CDRH3 lengths seen in *λ*-CSTs are due to a selection bias in therapeutic development.

**Fig. 4.**
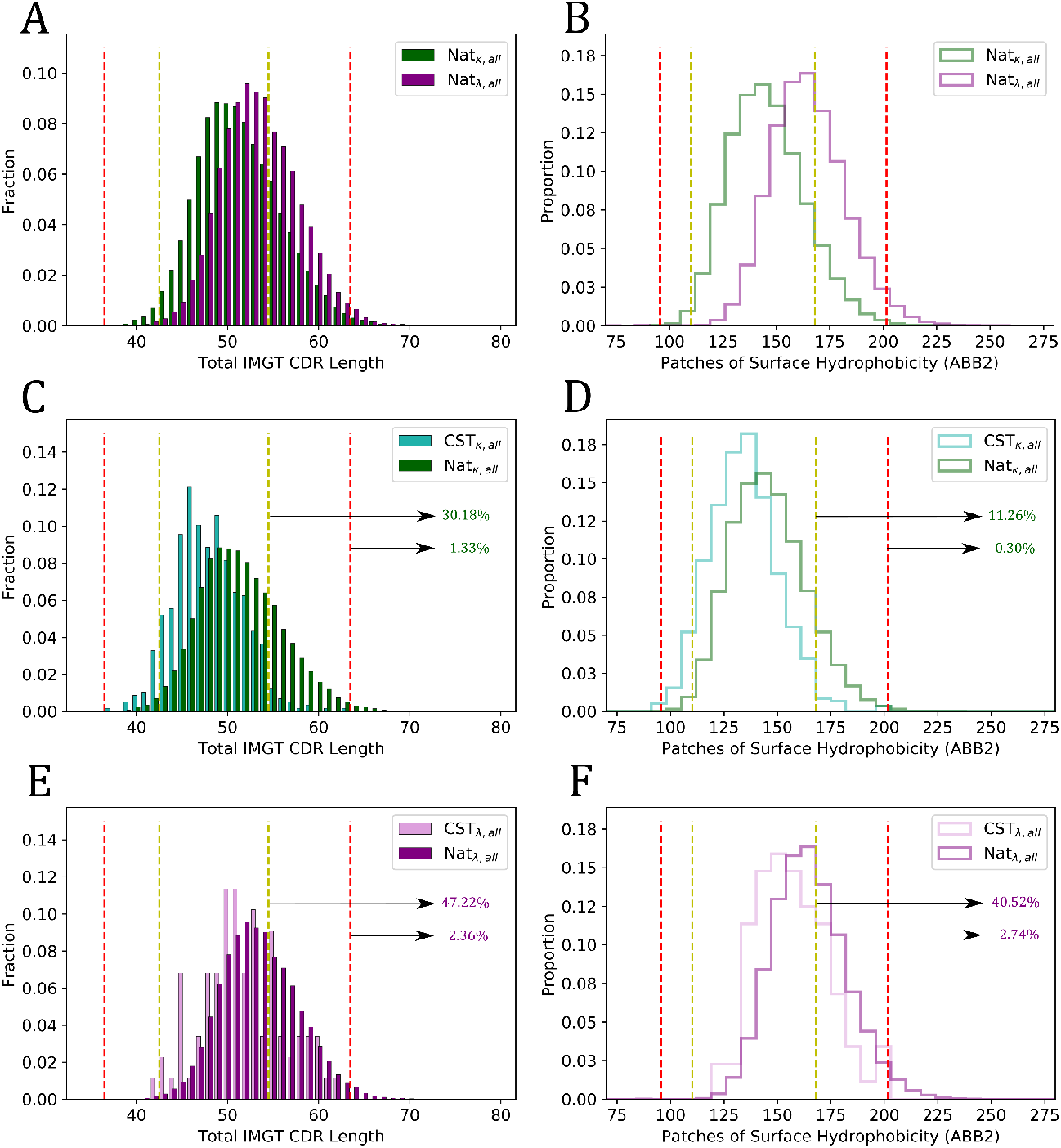
Plots of the Total CDR Length (L_tot_) and Patches of Surface Hydrophobicity (PSH) metric scores across different datasets split by light chain isotype. (A, B) L_tot_ and PSH for the *κ*-Nat_*all*_ and *λ*-Nat_*all*_ subsets. (C, D) L_tot_ and PSH for the *κ*-CST_all_ and *κ*-Nat_all_ subsets. (E, F) L_tot_ and PSH for the *λ*-CST_all_ and *λ*-Nat_all_ subsets. Highlighted percentages show the proportions of *κ*-Nat_all_ and *λ*-Nat_all_ antibodies exceeding the upper TAP thresholds. ABB2: ABodyBuilder2 models.

Analogous to the CSTs, natural human *λ* antibodies were disproportionately flagged for high PSH relative to human *κ*-antibodies, but both were flagged at an even higher rate: 11.26% of natural *κ*-antibodies and 40.52% of natural *λ-* antibodies flagged, relative to 1.91% and 26.14% of *κ*-CSTs and *λ*-CSTs, respectively.

Both *κ*-CSTs and *λ*-CSTs therefore occupy a lower-risk region of CDR length and PSH property space relative to natural antibodies, strengthening the findings from the original TAP paper (18) where we suggested that CSTs in general require more conservative values of these properties than natural antibodies to be amenable to therapeutic development. It also highlights the complexity of developability optimisation in drug discovery: improvements in the ‘humanness’ of the antibodies in screening libraries can have the unintended byproduct of enhanced therapeutic aggregation risk, regardless of the genetics of the light chain.

The charge properties of the natural *κ*- and *λ*-antibodies can be found in Fig. S10. The natural *λ*-antibodies also show an enhanced propensity for PPC values over 1 relative to their kappa equivalents, suggesting a natural origin for the disproportionate flagging of *λ*-CSTs for PPC (Fig. 3C).

### Residue positions associated with driving *λ*-antibodies towards high PSH scores

Our analyses suggest that the structure-dependent property biases across *λ*-CSTs are inherited from natural trends, especially for the PSH score. We therefore examined the *λ*-CSTs to determine which features in the Fv tend to correlate with their high PSH scores, with a view to guiding rational engineering and library design.

We selected the red-flagging sets of natural *κ*- and *λ*-antibodies and decomposed the overall TAP PSH score into its pairwise-residue component parts. We investigate in more detail the top-20-most hydrophobic sequence-adjacent interactions, and top-20-most hydrophobic sequence non-adjacent interactions, across antibodies red-flagging for PSH.

We observed a broad diversity of heavy (Fig. S11) and light chain (Fig. S12) residues involved in driving extreme PSH scores, emphasising the challenge of finding molecular rules of thumb for antibody optimisation. However, the heavy residue positions involved in elevating the PSH of either *κ*-antibodies or *λ*-antibodies were highly similar, suggesting minimal bias in the physicochemical properties of heavy chains associating with *κ*- or *λ*-antibodies.

Of particular interest to antibody optimisation engineers are positions outside of the CDR regions, since mutations at these sites are less likely to impact antigen specificity. Amongst the dominant residues contributing to high PSH scores were *κ*-VL positions 1-3 (framework region L1) and 79-85 (framework region L3), and *λ*-VL positions 24-26 (framework region L1) and 71-72 (framework region L3); while not in the formal CDRs these residues lie in the vicinity of the CDRs and may be serving to extend hydrophobic self-association surfaces.

As a case study, we investigated in more detail light chain positions 24-26, which drive higher PSH scores in *λ*-antibodies but not *κ*-antibodies (Fig. 5A, Fig. S13). The *λ*-antibodies exhibit a broader diversity of amino acids at these positions than *κ*-antibodies, although, with the exception of leucine at position 25 (Fig. S13), do not exhibit particularly hydrophobic residues. However, we observed that over 90% of *κ*-antibodies have positively-charged residues at position 24, while *λ*-antibodies almost exclusively use smaller, less polar residues (Fig. 5A), which would be expected to be more accommodating of a hydrophobic self-association interface. Meanwhile, though serine is the most commonly observed residue at position 26 in *λ*-antibodies (and seen in 100% of *κ*-antibodies, Fig. S13), threonine becomes by far the most prevalent residue amongst red-flagging *λ*-antibodies when position 26 is involved in the subset of most hydrophobic interactions (Fig. 5B). This is due both to its slightly higher intrinsic hydrophobicity and to more complex co-associations with other residues.

**Fig. 5.**
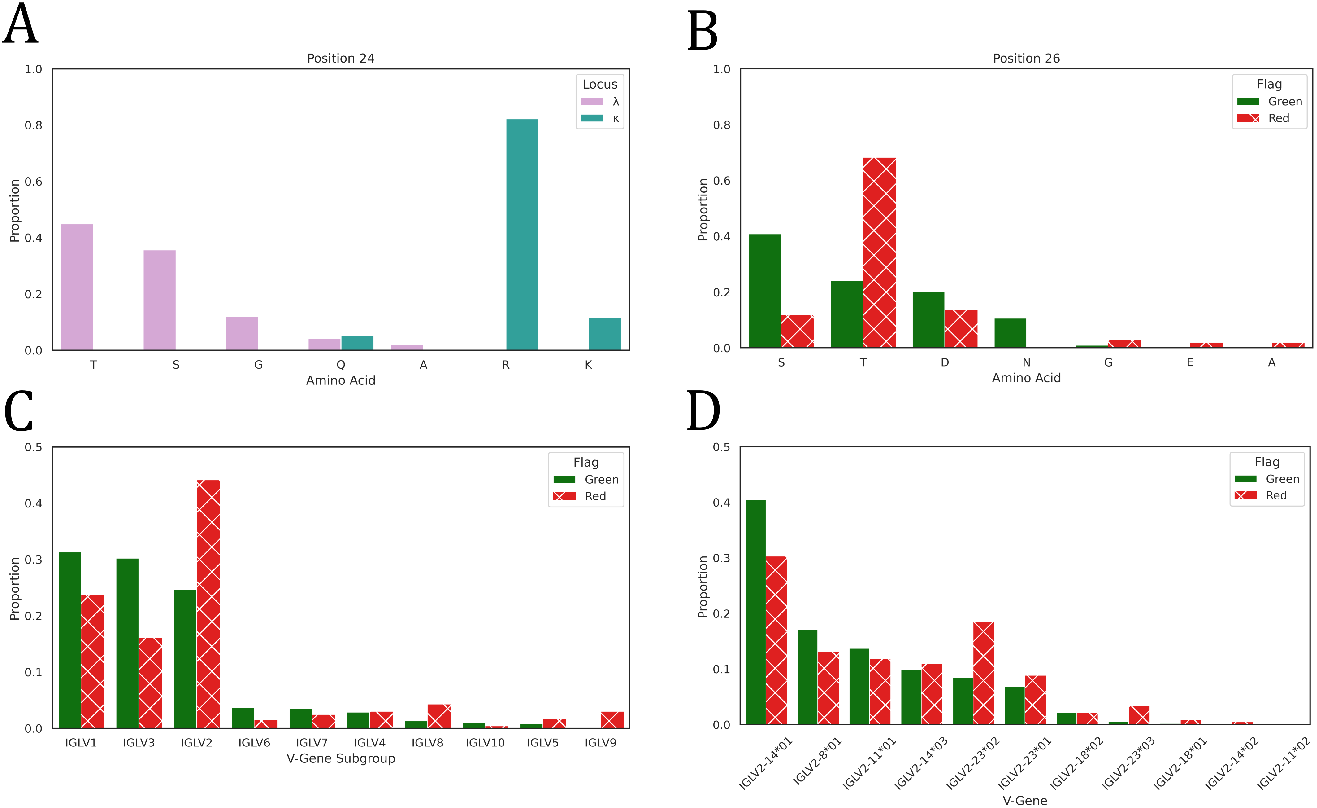
A: Bar charts showing the amino acid usages at IMGT position 24 amongst natural *λ*-antibodies (plum) and natural *κ*-antibodies (seagreen). B: Bar charts showing the amino acid usage at IMGT position 26 across natural *λ*-antibodies that are green-flagged for PSH, or that are red-flagged where position 26 is involved in the subset of most hydrophobic interactions. C-D: Bar charts showing (C) the light V gene subgroup usages amongst natural *λ*-antibodies that are green-flagged or red-flagged for PSH, and (D) the IGLV2 gene usages amongst natural *λ*-antibodies that are green-flagged or red-flagged for PSH.

In summary, by decomposing TAP scores such as the PSH into pairwise residue components, we can elucidate the regions driving high risk scores for individual or classes of antibodies and help to orient developability-motivated mutagenesis studies.

### *λ*-VL genes harbour characteristic risk profiles

Associations of certain genes or gene families with PSH flagging propensity would offer a simple strategy to develop diverse but developable screening libraries, for example by incorporating only select lambda genes with a more moderate risk of poor developability. To investigate if such associations exist, we evaluated the gene usages of *λ*-antibodies that green-flagged or red-flagged for PSH (Fig. 5C-D). From a gene family perspective (Fig. 5C), IGLV1 and IGLV3 were associated with a lower PSH-mediated developability risk, while others such as IGLV2, were highly enriched amongst red-flagging *λ*-antibodies. IGLV9 is almost exclusively found amongst flagging antibodies. These risk profiles are supported by the gene family usages across *λ*-CSTs (Table S8): IGLV1 and IGLV3 are over-represented relative to their natural abundances, while IGLV2 is under-represented, and no CST has yet derived from an IGLV9 gene.

To study what features might be driving differential risk across families, we generated separate sequence logo plots of all IGLV2 sequences and all non-IGLV2 sequences (Fig. S14). This highlighted positions that are commonly more hydrophobic among IGLV2 antibodies. For example, position 57 is mostly valine with a trace of glycine in IGLV2 antibodies, whereas is predominantly asparagine or asparatate in antibodies from the other families. Similarly, position 3 is entirely hydrophobic across IGLV2 antibodies but is found to be glutamate in roughly 1/3 of the other LV gene subgroups. Consistent with Fig. 5B, position 26 is almost entirely thre-onine in IGLV2 antibodies, while other families tend to use the less hydrophobic serine and highly polar residues such as asparagine and aspartate. Additionally, we observed that the CDRL1 loop, which typically bears a central motif containing hydrophobic residues, is frequently longer, and therefore more protruding, in IGLV2 antibodies.

We then dissected the TAP PSH risk profiles for the IGLV2 antibodies into profiles for individual genes (Fig. 5D). This demonstrated that the higher developability risk associated with the family is not shared evenly amongst its constituent genes: for example, IGLV2-14*01 is found in a higher fraction of green-flagging antibodies than red-flagging antibodies, while every allele of IGLV2-23 is associated with enhanced abundance among red-flagging *λ*-antibodies. Again, this is supported by gene usages across *λ*-CSTs: IGLV2-14 is the dominant gene observed amongst the relatively small number of CSTs deriving from the IGLV2 family (Table S9). Together these results suggest that a *λ*-antibody’s gene origin contributes substantially to its developability risk profile, and that TAP can be used to stratify lower from higher risk scaffolds.

## Discussion

In this paper, we benchmarked the latest machine learning-based antibody modeling technology for use in the Therapeutic Antibody Profiler (18).

We found that, while the precise guideline values we derived in 2019 have modulated slightly due to the availability of nearly three-times as many CST datapoints, the broad trends in property distribution between CSTs and natural antibodies have remained consistent; *i*.*e*. CSTs as a whole have significantly shorter CDR loops and smaller patches of surface hydrophobicity, while their charge properties are highly similar. The patterns also hold when limited to the subset of higher-certainty models (based on ABodyBuilder2’s statistical heuristic (30)).

When split by year of designation by the WHO, new therapeutics are more frequently sampling the extremes of CDR and PSH property space, indicating that our definitions of ‘druglikeness’ are likely to continue evolving over time. This phenomenon has also been observed in small molecule drug discovery, where several typical properties of today’s drugs differ from the original trends documented by Lipinski et al. (40, 41), and may be related to advances in developmental/formulation technologies. On the other hand, we observe no obvious trends in the properties of post-Phase I active/approved therapeutics *versus* discontinued therapeutics, nor in therapeutics that have advanced to different clinical stages, suggesting that, at least in terms of the TAP properties, we would not expect predictive power to improve by only considering therapeutics that have advanced to later stages.

Due to ABodyBuilder2’s modeling protocol, statistical uncertainty in side chain positioning can now be captured to some extent through repeat modeling and TAP calculations. Guidelines derived from repeat runs are almost identical to the guidelines derived from a single run per therapeutic, while mean variances of property values of the CST therapeutics are near-0 for charge metrics and only around 10 for the PSH metric. Variances on this order can lead to classification disparities across repeat runs between adjacent boundaries (*i*.*e*. green/amber risk, or amber/red risk) but are extremely unlikely to lead to the same antibody being assigned green risk and red risk for a given property.

Molecular dynamics simulations of a representative set of CST Fab models indicate that the flags assigned by an ensemble of repeat static ABodyBuilder2 predictions are highly consistent with simulation-average flags. Best agreement with simulation is obtained by considering an antibody to have flagged for a property if a flag is seen on any of the repeat runs. As running TAP multiple times takes a few minutes, several orders of magnitude faster than running molecular dynamics, repeat TAP calculations may offer a sensible strategy for high-throughput developability screening with consideration for side chain mobility.

We then used our new TAP protocol to investigate developability-linked property biases across *κ*- and *λ*-antibodies, exploiting the rise in both CST and paired-chain natural sequence data. *λ*-VLs have distinct epitope specificities to *κ*-VLs, driven by features such as locus-specific germline-encoded amino acid binding-motifs (10). However *λ*-antibodies have been anecdotally found to be less developable than *κ*-antibodies (25) and are heavily under-sampled amongst CSTs relative to their natural abundance. Therapeutic antibody profiling adds quantitative evidence that natural *λ*-antibodies are generally at higher risk of developablity issues, especially hydrophobicity-driven aggregation, than natural *κ*-antibodies. Indeed, the mean of the natural *λ*-antibody distribution sits just below the amber-flagging threshold; a substantial population of *λ*-antibodies are prone to being nudged into being flagged by, say, an affinity maturation pipeline based on unconstrained mutagenesis.

However, through a quantification of the risk of each *λ-*antibody, TAP profiles can now enable us to identify subpopulations expected to be more amenable to therapeutic development, and therefore to offer strategies towards augmenting the targetable epitope space through rational design. The observation of particular lambda gene associations with higher risk profiles, and a preliminary concomitant signal in the gene origin distributions of *λ*-CSTs that have so far progressed to the clinic, suggest that approaches such as family-holdout (e.g. all IGLV2) or gene-holdout (e.g. all IGLV2-23) libraries should enrich for *λ*-antibodies with lower expected developability risk. Alternatively, libraries could be constructed at a more granular level, incorporating more risk-prone genes but only when the associated sequence is considered by TAP to be lower risk. While sequence-by-sequence *in vitro* screening library design may still be a distant prospect, such approaches are gaining traction in the field of *in silico* library design (42, 43).

The interpretability of the TAP metrics means they can be readily deconstructed to explore which regions of the CDR vicinity tend to contribute to higher developability risk. We show that the positions that contribute most to high risk scores in both *κ*- and *λ*-antibodies are diverse and distinct. Differences lie in the light chain itself rather than through any biases in the properties of their paired heavy chains. While preliminary, we note that some residues in the periphery of the CDR vicinity can help drive antibodies towards being red flagged; on a case-by-case basis, mutations to these regions may impact developability while being less likely to affect antibody specificity.

To date, TAP has primarily found use in industry for the early-stage removal of candidate antibodies more likely to suffer from developability issues. This increases the efficiency of drug discovery, but risks artificially constraining diversity, reinforcing current established property trends. We have shown how TAP, applied to identify more nuanced *λ*-

VL residue and *λ*-gene associations with developability risk, could also guide selective consideration of a broader diversity of lead candidates and so enable access to a wider pool of epitopes.

## Methods

### Dataset Curation

The Therapeutic Structural Antibody Database (Thera-SAbDab) was downloaded on 25^th^ January, 2023 (1). The entries were filtered for those designed for human application, that have reached at least Phase-II of clinical trials, and that have complete variable regions (Fvs, *i*.*e*. no single domain antibodies were carried forward). This set was then mined for sequence non-redundant Fv regions (at the level of 100% identity), to filter out biosimilars with no changes to the Fv and to reduce biases caused by the use of previously-developed monoclonal Fv domain sequences in new multispecific formats. This resulted in 664 non-redundant CST Fvs (the CST_all_ dataset), of which 576 were *κ*-based (86.7%) and 88 were *λ*-based (13.3%). Thera-SAbDab light-gene locus labels were confirmed *via* alignment to the latest set of human and mouse IMGT V domains using ANARCI (44).

The sequence non-redundant (100% identity) Fv sequences of 88,274 natural human antibodies were retrieved from the Observed Antibody Space (OAS) database (29) (timestamp: 25^th^ January 2023). These were filtered for sequences with complete CDRs (45), leaving 79,761 antibodies (the Nat_all_ dataset): 44,420 (55.7%) *κ*-antibodies, 35,341 (44.3%) *λ*-antibodies.

### Benchmarking TAP Modeling Methods

ABodyBuilder1 (31) was run using template databases built on a copy of SAbDab (45, 46) timestamped to 30^th^ April 2022, and with a template sequence identity cut-off of 99% to ensure genuine models were produced (31). ABodyBuilder2 was run using the pre-trained weights from the paper (34). Relative performance to ABodyBuilder1 was evaluated by root-mean-squared deviation and the percentage of side chain residues correctly classified as surface exposed (see Supporting Information Methods).

A threshold to filter out the least confident ABodyBuilder2 models was obtained by a two-step process. First, the 119/664 CST Fv domains for which 100% sequence iden-tical X-ray crystal structures exist (identified using Thera-SAbDab metadata (1), Table S2), were filtered out of the CST_all_ dataset. The root-mean-squared predicted error for each remaining CST CDRH3 was then calculated 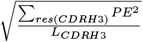, where PE represents backbone predicted error, res(CDRH3) represents the sum over all CDRH3 residues and L(CDRH3) represents the length of the CDRH3. Finally, the threshold was derived by evaluating the 75^th^ percentile (1.31 Å). This filter was applied to retain the 510 most confidently-modeled CSTs (the CST_conf_ dataset), and applied to the Nat_all_ dataset to derive the 30,402 most confidently modeled natural antibodies (the Nat_conf_ subset).

### Running TAP on ABodyBuilder2 models

Sets of CST and natural antibody Fv domains were run through TAP and their five computational developability metrics calculated (Total IMGT CDR Length [L_tot_], Patches of Surface Hydrophobicity using the Kyte and Doolittle scale [PSH], Patches of Surface Positive Charge [PPC], Patches of Surface Negative Charge [PNC] and Structural Fv Charge Symmetry Parameter [SFvCSP]) (18). PSH, PPC, and PNC metrics were calculated across the CDR vicinity (IMGT-defined CDR residues *±* 2 on each side plus any other surface exposed residue within 4.5 Å of one of these residues). Throughout the work, amber and red thresholds and were set at the percentile values suggested in the original paper (18).

Whenever the properties of *κ*- and *λ*-antibodies where compared, threshold values were calculated from the CST_all_ set (*i*.*e*. not evaluated separately by light chain type).

### Assessing TAP score variation over molecular dynamics trajectories

14 CST Fab regions were modelled by grafting the constant regions of their crystal structures (see Table S6) onto the Fv models generated by ABodyBuilder2, obtaining the initial arrangement by aligning the crystal and model Fv backbones (full Fab regions were used instead of Fv regions based on the results of previous studies (47)). We then modelled-in missing residues in the constant region using MODELLER v10.2 (48) and generated 10 models using the ‘very slow’ refinement setting, selecting the lowest energy model. All systems were prepared and simulations performed using OpenMM v7.7 (49). N-methyl groups were used to cap C-termini using an in-house script. Next, using ‘pdbfixer’ (49), we protonated the models at a pH of 7.5, soaked them in truncated octahedral water boxes with a padding distance of 1 nm, and added sodium or chloride counter-ions to neutralise charges and then NaCl to an ionic strength of 150 mM. We parameterised the systems using the Amber14-SB forcefield (50) and modelled water molecules using the TIP3P-FB model (51). Non-bonded interactions were calculated using the particle mesh Ewald method (52) using a cut-off of distance of 0.9 nm, with an error tolerance of 5×10^−4^. Water molecules and heavy atom-hydrogen bonds were rigidified using the SETTLE (53) and SHAKE (54) algorithms, respectively. We used hydrogen mass repartitioning (55) to allow for 4 fs time steps. Simulations were run using the mixed-precision CUDA platform in OpenMM us-ing the Middle Langevin Integrator with a friction coefficient of 1 ps^-1^ and the Monte-Carlo Barostat set to 1 atm. We equilibrated systems using a multi-step protocol detailed in Table S7. Following equilibration, we performed 200 ns of unrestrained simulation of the NPT ensemble at 300K, calculating TAP properties over the final 120 ns of each simulation, when all systems had reached relatively stable RMSD values from their initial coordinates (Fig. S15). To estimate convergence, we aligned Fv regions on the starting structure using mdtraj v1.9.6 (56) and calculated the RMSD of the Fv domains relative to the starting structure.

### Determining molecular correlates with poor developability

Natural human antibodies lying above the red flag thresholds set by TAP across all CSTs were investigated for recurrent molecular patterns that contribute towards their high scores. The PSH scores for each antibody were split into components from sequence-adjacent residues and components from sequence non-adjacent residues, and these pairwise interactions were separately rank-ordered by hydrophobicity. Germline assignments for natural sequences were taken from the OAS Paired metadata (29), which derives from IgBlast (57) alignments of each nucleotide sequence to a recent set of human genes from the IMGT GeneDB (4). Germline assignments for CSTs were evaluated using AN-ARCI (44) on amino acid sequences (allele predictions were ignored here due to the difficulty of accurately assigning alleles at the amino acid level). All percentage abundances of gene/gene family usages across *λ*-CSTs were calculated based the subset that mapped closest to human rather than mouse germlines.

### Visualisations

All visualisations were made using open-source PyMOL or matplotlib version 3.5.2.

## Code and Data Availability

The updated TAP protocol is available on our web application (https://opig.stats.ox.ac.uk/webapps/tap/) and through the Vagrant Virtual Machine and Singularity container versions of our SAbDab-SAbPred codebase. The structures used for ABodyBuilder2 benchmarking are available as Dataset S1. The ABodyBuilder2 models of all CSTs analysed in this study are available as Dataset S2. ABodyBuilder2 models of natural paired-chain human antibodies were released in the Supporting Materials of Abanades *et al*. (30).

## Acknowledgements

The authors would like to thank Dr. Sandeep Kumar (Boehringer Ingelheim) for critically reviewing the manuscript. This work was supported by a Postdoctoral Research grant funded by Boehringer Ingelheim (MR), funding from the UK Engineering and Physical Sciences Research Council (OT, reference EP/S024093/1), and the Wellcome Trust (BG, reference 102164/Z/13/Z).

## Bibliography

1. Matthew I J Raybould, Claire Marks, Alan P Lewis, Jiye Shi, Alexander Bujotzek, Bruck Taddese, and Charlotte M Deane. Thera-SAbDab: the Therapeutic Structural Antibody Database. Nucleic Acids Res, 48(D1):D383–D388, 2019. doi: 10.1093/nar/gkz827.

2. Melanie M. Senior. Fresh from the biotech pipeline: fewer approvals, but biologics gain share. Nat Biotechnol, 42(2):174–182, 2023. doi: 10.1038/s41587-022-01630-6.

3. Asher Mullard. FDA approves 100th monoclonal antibody product. Nat Rev Drug Discov, 20:491–495, 2021. doi: 10.1038/d41573-021-00079-7.

4. Véronique Giudicelli, Denys Chaume, and Marie-Paule Lefranc. IMGT/GENE-DB: a comprehensive database for human and mouse immunoglobulin and T cell receptor genes. Nucleic Acids Res, 33(D1):D256–D261, 2005. doi: doi:10.1093/nar/gki010.

5. Anthony R. Rees. Understanding the human antibody repertoire. mAbs, 12(1):1729683, 2020. doi: 10.1080/19420862.2020.1729683.

6. Kenneth Smith, Hemangi Shah, Jennifer J. Muther, Angie L. Duke, Kathleen Haley, and Judith A. James. Antigen nature and complexity influence human antibody light chain usage and specificity. Vaccine, 34(25):2813–2820, 2016. doi: https://doi.org/10.1016/j.vaccine.2016.04.040.

7. Matthew I. J. Raybould, Aleksandr Kovaltsuk, Claire Marks, and Charlotte M. Deane. CoV-AbDab: the coronavirus antibody database. Bioinformatics, 37(5):734–735, 2021. doi: 10.1093/bioinformatics/btaa739.

8. Sarah A. Robinson, Matthew I. J. Raybould, Constantin Schneider, Wing Ki Wong, Claire Marks, and Charlotte M. Deane. Epitope profiling using computational structural modelling demonstrated on coronavirus-binding antibodies. PLoS Comput Biol, 17(12):e1009675, 2021. doi: 10.1371/journal.pcbi.1009675.

9. David B. Jaffe, Payam Shahi, Bruce A. Adams, Ashley M. Chrisman, Peter M. Finnegan, Nandhini Raman, Ariel E. Royall, FuNien Tsai, Thomas Vollbrecht, Daniel S. Reyes, and Wyatt J. McDonnell. Functional antibodies exhibit light chain coherence. Nature, 611:352–357, 2022. doi: 10.1038/s41586-022-05371-z.

10. Ellen L. Shrock, Richard T. Timms, Tomasz Kula, Elijah L. Mena, Anthony P. West Jr., Rui Guo, I-Hsiu Lee, Alexander A. Cohen, Lindsay G. A. McKay, Caihong Bi Keerti, Yumei Leng, Eric Fujimara, Felix Horns, Mamie Li, Duane R. Wesemann, Anthony Griffiths, Benjamin E. Gewurz, Pamela J. Bjorkman, and Stephen J. Elledge. Germline-encoded amino acid–binding motifs drive immunodominant public antibody responses. Science, 380(6640), 2023. doi: 10.1126/science.adc94.

11. Brandon J. DeKosky, Oana I. Lungu, Daechan Park, Erik L. Johnson, Wissam Charab, Constantine Chrysostomou, Daisuke Kuroda, Andrew D. Ellington, Gregory C. Ippolito, Jeffrey J. Gray, and George Georgiou. Large-scale sequence and structural comparisons of human naive and antigen-experienced antibody repertoires. Proc Natl Acad Sci, 113(19): E2636–E2645, 2016. doi: 10.1073/pnas.1525510113.

12. Puneet Rawat, R. Prabakaran, Sandeep Kumar, and M. Michael Gromiha. Exploring the sequence features determining amyloidosis in human antibody light chains. Sci Rep, 11: 13785, 2021. doi: 10.1038/s41598-021-93019-9.

13. William S. Gibson, Oscar L. Rodriguez, Kaitlyn Shields, Catherine A. Silver, Abdullah Dorgham, Matthew Emery, Gintaras Deikus, Robert Sebra, Evan E. Eichler, Ali Bashir, Melissa L. Smith, and Corey T. Watson. Characterization of the immunoglobulin lambda chain locus from diverse populations reveals extensive genetic variation. Genes Immun, 24:21–31, 2023. doi: 10.1038/s41435-022-00188-2.

14. Robyn L. Stanfield, Adam Zemla, Ian A. Wilson, and Bernhard Rupp. Antibody Elbow Angles are Influenced by their Light Chain Class. J Mol Biol, 357(5):1566–1574, 2006. doi: 10.1016/j.jmb.2006.01.023.

15. Daisuke Kuroda, Hiroki Shirai, Masato Kobori, and Haruki Nakamura. Systematic classification of CDR-L3 in antibodies: implications of the light chain subtypes and the VL-VH interface. Proteins, 75(1):139–146, 2009. doi: doi:10.1002/prot.22230.

16. Rob van der Kant, Joschka Bauer, Anne R Karow-Zwick, Sebastian Kube, Patrick Garidel, Michaela Blech, Frederic Rousseau, and Joost Schymkowitz. Adaption of human antibody λ and κ light chain architectures to CDR repertoires. Protein Engineering, Design and Selection, 32(3):109–127, 2019. doi: 10.1093/protein/gzz012.

17. Catherine L. Townsend, Jule M. J. Laffy, Yu-Chang B. Wu, Silva O’Hare, Victoria Martin, David Kipling, Franca Fraternali, and Deborah K. Dunn-Walters. Significant Differences in Physicochemical Properties of Human Immunoglobulin Kappa and Lambda CDR3 Regions. Front Immunol, 7:388, 2016. doi: 10.3389/fimmu.2016.00388.

18. Matthew I. J. Raybould, Claire Marks, Konrad Krawczyk, Bruck Taddese, Jaroslaw Nowak, Alan P. Lewis, Alexander Bujotzek, Jiye Shi, and Charlotte M. Deane. Five computational developability guidelines for therapeutic antibody profiling. Proc Natl Acad Sci, 116(10): 4025–4030, 2019. doi: 10.1073/pnas.1810576116.

19. Claire M. Molé, Marie C. Béné, Paul M. Montagne, Estelle Seilles, and Gilbert C. Faurea. Light chains of immunoglobulins in human secretions. Clin Clim Acta, 224(2):191–197, 1994. doi: 10.1016/0009-8981(94)90185-6.

20. Aleksandr Kovaltsuk, Jinwoo Leem, Sebastian Kelm, James Snowden, Charlotte M. Deane, and Konrad Krawczyk. Observed Antibody Space: A Resource for Data Mining Next-Generation Sequencing of Antibody Repertoires. J Immunol, 201(8):2502–2509, 2018. doi: 10.4049/jimmunol.1800708.

21. Ruei-Min lu, Yu-Chyi Hwang, I-Ju Liu, Chi-Chiu Lee, Han-Zen Tsai, Hsin-Jung Li, and Han-Chung Wu. Development of therapeutic antibodies for the treatment of diseases. J Biomed Sci, 27:1, 2020. doi: 10.1186/s12929-019-0592-z.

22. Andreas H. Laustsen, Victor Greiff, Aneesh Karatt-Vellatt, Serge Muyldermans, and Timothy P. Jenkins. Animal Immunization, in vitro Display Technologies, and Machine Learning for Antibody Discovery. Trends Biotechnol, 39(12):1263–1273, 2021. doi: 10.1016/j.tibtech.2021.03.003.

23. Andre Azevedo Reis Teixeira, Michael Frank Erasmus, Sara D’Angelo, Leslie Naranjo, Fortunato Ferrara, Camila Leal-Lopes, Oliver Durrant, Cecile Galmiche, Aleardo Morelli, Anthony Scott-Tucker, and Andrew Raymon Morton Bradbury. Drug-like antibodies with high affinity, diversity and developability directly from next-generation antibody libraries. mAbs, 13(1):1980942, 2021. doi: 10.1080/19420862.2021.1980942.

24. Mani Larijani, Shuang Chen, Lesley A. Cunningham, Joseph M. Volpe, Lindsay Grey Cowell, Susanna M. Lewis, and Gillian E. Wu. The recombination difference between mouse kappa and lambda segments is mediated by a pair-wise regulation mechanism. Mol Immunol, 43 (7):870–881, 2006. doi: 10.1016/j.molimm.2005.06.038.

25. Andreas Lehmann, Josephine H. F. Wixted, Maxim V. Shapovalov, Heinrich Roder, Roland L. Dunbrack Jr., and Matthew K. Robinson. Stability engineering of anti-EGFR scFv antibodies by rational design of a lambda-to-kappa swap of the VL framework using a structure-guided approach. mAbs, 7(9):1058–1071, 2015. doi: 10.1080/19420862.2015.1088618.

26. Kip Bodi, Tatiana Prokaeva, Brian Spencer, Maurya Eberhard, Lawreen H. Connors, and David C. Seldin. AL-Base: a visual platform analysis tool for the study of amyloidogenic immunoglobulin light chain sequences. Amyloid, 16(1):1–8, 2009. doi: 10.1080/13506120802676781.

27. Juan C. Almagro, Martha Pedraza-Escalon, Hugo I. Arrieta, and Sonia M. Pérez-Tapia. Phage Display Libraries for Antibody Therapeutic Discovery and Development. Antibodies, 8(3):44, 2019. doi: 10.3390/antib8030044.

28. Sandeep Kumar, Kirk Roffi, Dheeraj Tomar, David Cirelli, Nicholas Luksha, Danielle Meyer, Jeffrey Mitchell, Martin J. Allen, and Li Li. Rational optimization of a monoclonal antibody for simultaneous improvements in its solution properties and biological activity. Prot Eng Des Sel, 31(7-8):313–325, 2018. doi: 10.1093/protein/gzy020.

29. Tobias H. Olsen, Fergus Boyles, and Charlotte M. Deane. Observed Antibody Space: A diverse database of cleaned, annotated, and translated unpaired and paired antibody sequences. Protein Sci, 31(1):141–146, 2022. doi: 10.1002/pro.4205.

30. Brennan Abanades, Wing Ki Wong, Fergus Boyles, Guy Georges, Alexander Bujotzek, and Charlotte M. Deane. ImmuneBuilder: Deep-Learning models for predicting the structures of immune proteins. Commun Biol, 6:575, 2023. doi: 110.1038/s42003-023-04927-7.

31. Jinwoo Leem, James Dunbar, Guy Georges, Jiye Shi, and Charlotte M. Deane. ABody-Builder: Automated antibody structure prediction with data–driven accuracy estimation. mAbs, 8(7):1259–1268, 2016. doi: 10.1080/19420862.2016.1205773.

32. Richard Evans, Michael O’Neill, Alexander Pritzel, Natasha Antropova, Andrew Senior, Tim Green, Augustin Zídek, Russ Bates, Sam Blackwell, Jason Yim, Olaf Ronneberger, Sebastian Bodenstein, Michal Zielinski, Alex Bridgland, Anna Potapenko, Andrew Cowie, Kathryn Tunyasuvunakool, Rishub Jain, Ellen Clancy, Pushmeet Kohli, John Jumper, and Demis Hassabis. Protein complex prediction with AlphaFold-Multimer. bioRxiv, 2022. doi: 10.1101/2021.10.04.463034.

33. Jeffrey A. Ruffolo, Lee-Shin Chu, Sai Pooja Mahajan, and Jeffrey J. Gray. Fast, accurate antibody structure prediction from deep learning on massive set of natural antibodies. Nat Commun, 14:2389, 2023. doi: 10.1038/s41467-023-38063-x.

34. Brennan Abanades, Guy Georges, Alexander Bujotzek, and Charlotte M Deane. ABlooper: fast accurate antibody CDR loop structure prediction with accuracy estimation. Bioinformatics, 38(7):1877–1880, 2022. doi: 10.1093/bioinformatics/btac016.

35. Giuseppe Licari, Kyle P. Martin, Maureen Crames, Joseph Mozdzierz, Michael S. Marlow, Anne R. Karow-Zwick, Sandeep Kumar, and Joschka Bauer. Embedding Dynamics in Intrinsic Physicochemical Profiles of Market-Stage Antibody-Based Biotherapeutics. Mol Pharmaceutics, 20(2):1096–1111, 2023. doi: 10.1021/acs.molpharmaceut.2c00838.

36. Lucky Ahmed, Priyanka Gupta, Kyle Martin, Justin M. Scheer, Andrew M. Nixon, and Sandeep Kumar. Intrinsic physicochemical profile of marketed antibody-based biotherapeutics. Proc Natl Acad Sci, 118(37):e2020577118, 2021. doi: 10.1073/pnas.2020577118.

37. Leonard D. Goldstein, Ying-Jiun J. Chen, Jia Wu, Subhra Chaudhuri, Yi-Chun Hsiao, Kellen Schneider, Kam Hon Hoi, Zhonghua Lin, Steve Guerrero, Bijay S. Jaiswal, Jeremy Stinson, Aju Antony, Kanika Bajaj Pahuja, Dhaya Seshasayee, Zora Modrusan, Isidro Hötzel, and Somasekar Seshagiri. Massively parallel single-cell B-cell receptor sequencing enables rapid discovery of diverse antigen-reactive antibodies. Commun Biol, 2:304, 2019. doi: 10.1038/s42003-019-0551-y.

38. Matthew I. J. Raybould, Claire Marks, Aleksandr Kovaltsuk, Alan P. Lewis, Jiye Shi, and Charlotte M. Deane. Public Baseline and shared response structures support the theory of antibody repertoire functional commonality. PLoS Comput Biol, 17(3):e1008781, 2021. doi: 10.1371/journal.pcbi.1008781.

39. wwPDB Consortium. Protein Data Bank: the single global archive for 3D macromolecular structure data. Nucleic Acids Res, 47(D1):D520–D528, 2019. doi: 10.1093/nar/gky949.

40. Christopher A. Lipinski, Franco Lombardo, Beryl W. Dominy, and Paul J. Feeney. Experimental and computational approaches to estimate solubility and permeability in drug discovery and development settings. Adv Drug Deliv Rev, 23(1–3):3–25, 1997. doi: 10.1016/S0169-409X(96)00423-1.

41. Ingo V. Hartung, Bayard R. Huck, and Alejandro Crespo. Rules were made to be broken. Nat Rev Chem, 7:3–4, 2023. doi: 10.1038/s41570-022-00451-0.

42. Tileli Amimeur, Jeremy M. Shaver, Randal R. Ketchem, J. Alex Taylor, Rutilio H. Clark, John Smith, Danielle Van Citters, Christine C. Siska, Pauline Smidt, Megan Sprague, Bruce A. Kerwin, and Dean Pettit. Designing Feature-Controlled Humanoid Antibody Discovery Libraries Using Generative Adversarial Networks. bioRxiv, 2020. doi: 10.1101/2020.04.12.024844.

43. Matthew I. J. Raybould, Claire Marks, Aleksandr Kovaltsuk, Alan P. Lewis, Jiye Shi, and Charlotte M. Deane. Public Baseline and shared response structures support the theory of antibody repertoire functional commonality. PLoS Comput Biol, 17(3):e1008781, 2021. doi: 10.1371/journal.pcbi.1008781.

44. James Dunbar and Charlotte M. Deane. ANARCI: antigen receptor numbering and receptor classification. Bioinformatics, 32(2):298–300, 2016. doi: 10.1093/bioinformatics/btv552.

45. James Dunbar, Konrad Krawczyk, Jinwoo Leem, Terry Baker, Angelika Fuchs, Guy Georges, Jiye Shi, and Charlotte M. Deane. SAbDab: the structural antibody database. Nucleic Acids Res, 42(D1):D1140–D1146, 2014. doi: 10.1093/nar/gkt1043.

46. Constantin Schneider, Matthew I. J. Raybould, and Charlotte M. Deane. SAbDab in the age of biotherapeutics: updates including SAbDab-nano, the nanobody structure tracker. Nucleic Acids Res, 50(D1):D1368–D1372, 2022. doi: 10.1093/nar/gkab1050.

47. Bernhard Knapp, James Dunbar, Marta Alcala, and Charlotte M. Deane. Variable Regions of Antibodies and T-Cell Receptors May Not Be Sufficient in Molecular Simulations Investigating Binding. J Chem Theory Comput, 13(7):3097–3105, 2017. doi: 10.1021/acs.jctc.7b00080.

48. Benjamin Webb and Andrej Sali. Comparative protein structure modeling using modeller. Curr Protoc Bioinformatics, 54(1):5.7.1–5.7.37, 2016. doi: 10.1002/cpbi.3.

49. Peter Eastman, Jason Swails, John D. Chodera, Robert T. McGibbon, Tu-tong Zhao, Kyle A. Beauchamp, Lee Ping Wang, Andrew C. Simmonett, Matthew P. Harrigan, Chaya D. Stern, Rafal P. Wiewiora, Bernard R. Brooks, and Vijay S. Pande. OpenMM 7: Rapid development of high per-formance algorithms for molecular dynamics. PLoS Comput Biol, 13(7): e1005659, 2017. doi: 10.1371/journal.pcbi.1005659.

50. James A. Maier, Carmenza Martinez, Koushik Kasavajhala, Lauren Wick-strom, Kevin E. Hauser, and Carlos Simmerling. ff14sb: Improving the accuracy of protein side chain and backbone parameters from ff99sb. J Chem Theory Comput, 11(8):3696–3713, 2015. doi: 10.1021/acs.jctc.5b00255.

51. Eyal Neria, Stefan Fischer, and Martin Karplus. Simulation of activation free energies in molecular systems. J Chem Phys, 105(5):1902–1921, 1996. doi: 10.1063/1.472061.

52. Tom Darden, Darrin York, and Lee Pedersen. Particle mesh Ewald: AnNlog(N) method for Ewald sums in large systems. J Chem Phys, 98(12):10089–10092, 1993. doi: 10.1063/1.464397.

53. Shuichi Miyamoto and Peter A. Kollman. Settle: An analytical version of the SHAKE and RATTLE algorithm for rigid water models. J Comp Chem, 13(8):952–962, 1992. doi: 10.1002/jcc.540130805.

54. Jean Paul Ryckaert, Giovanni Ciccotti, and Herman J. C. Berendsen. Numerical integration of the cartesian equations of motion of a system with constraints: molecular dynamics of n-alkanes. J Chem Phys, 23(3):327–341, 1977. doi: 10.1016/0021-9991(77)90098-5.

55. Chad W. Hopkins, Scott Le Grand, Ross C. Walker, and Adrian E. Roitberg. Long-time-step molecular dynamics through hydrogen mass repartitioning. J Chem Theory Comput, 11(4): 1864–1874, 2015. doi: 10.1021/ct5010406.

56. Robert T. McGibbon, Kyle A. Beauchamp, Matthew P. Harrigan, Christoph Klein, Jason M. Swails, Carlos X. Hernández, Christian R. Schwantes, Lee Ping Wang, Thomas J. Lane, and Vijay S. Pande. MDTraj: A Modern Open Library for the Analysis of Molecular Dynamics Trajectories. Biophys J, 109(8):1528–1532, 2015. doi: 10.1016/j.bpj.2015.08.015.

57. Jian Ye, Ning Ma, Thomas L. Madden, and James M. Ostell. IgBLAST: an immunoglobulin variable domain sequence analysis tool. Nucleic Acids Res, 41(W1):W34–W40, 2013. doi: 10.1093/nar/gkt382.

