## Supplementary Information for "Contextualising the developability risk of antibodies with lambda light chains using enhanced therapeutic antibody profiling"

#### SI Results

**Evaluating ABodyBuilder2's Performance on Unseen Therapeutics.** To ensure that ABodyBuilder2's improved general performance translates to clinical-stage therapeutics (CSTs), we mined Thera-SAbDab to identify eight CST variable regions (Fvs) whose first sequence identical X-ray crystal structures were released after 31$^{st}$ July 2021 (Table S1); the 119 CST Fvs with crystal structures released before this date (Table S2) would not provide a fair indication of expected performance as they would have formed part of the ABodyBuilder2 training set.

Evaluating the performance of ABodyBuilder2 and ABodyBuilder1 over these eight 'unseen' CSTs we found that ABodyBuilder2's accuracy ($\mu_{\mathrm{CDRH3}}$: 2.68 Å) was markedly better than ABodyBuilder1's ($\mu_{\mathrm{CDRH3}}$: 3.32 Å; see Table S1 for all CDRs). We also observed that ABodyBuilder2's higher backbone prediction accuracy across this subset of CSTs translated into an improved accuracy in the proportions of side chains assigned as buried/exposed, a key parameter in evaluating the structure-dependent TAP metric values (ABodyBuilder2: 96.39% accuracy, ABodyBuilder1: 95.99% accuracy, Table S1). Improvement was further magnified when considering only the formal IMGT CDR (1) residues (Table S1). Together with the results from the ImmuneBuilder preprint (2), this evidence motivated a change in the TAP protocol to use ABodyBuilder2 for structural modeling in place of ABodyBuilder1.

#### SI Methods

**Benchmarking Model Quality.** Root-mean-squared deviation (RMSD) by IMGT-defined region (1) was calculated with an in-house script that first aligns each model structure to the ground truth structure based on the backbone atoms of the framework region of the investigated chain and then calculates the RMSD over the backbone atoms of the residues of the region (for heavy or light chains in the IMGT numbering scheme, CDR1: residues 27-38, CDR2: 56-65, CDR3: 105-117).

The classification of residues as solvent exposed or buried was based on an in-house implementation of the Shrake and Rupley algorithm (3), using a spherical probe of radius 1.4 Å. A residue 'X' was considered exposed if its solvent-accessible surface area (SASA) was $\geq$ 7.5% of its theoretical maximum value (based on the open-chain form of Alanine-X-Alanine) (4). In accordance with the parametrisations of these theoretical maximum SASAs, all hydrogen atoms were stripped out of ABodyBuilder2 predictions prior to SASA calculations.

#### SI Figures and Tables

The following pages contain **15** SI Figures and **9** SI Tables.

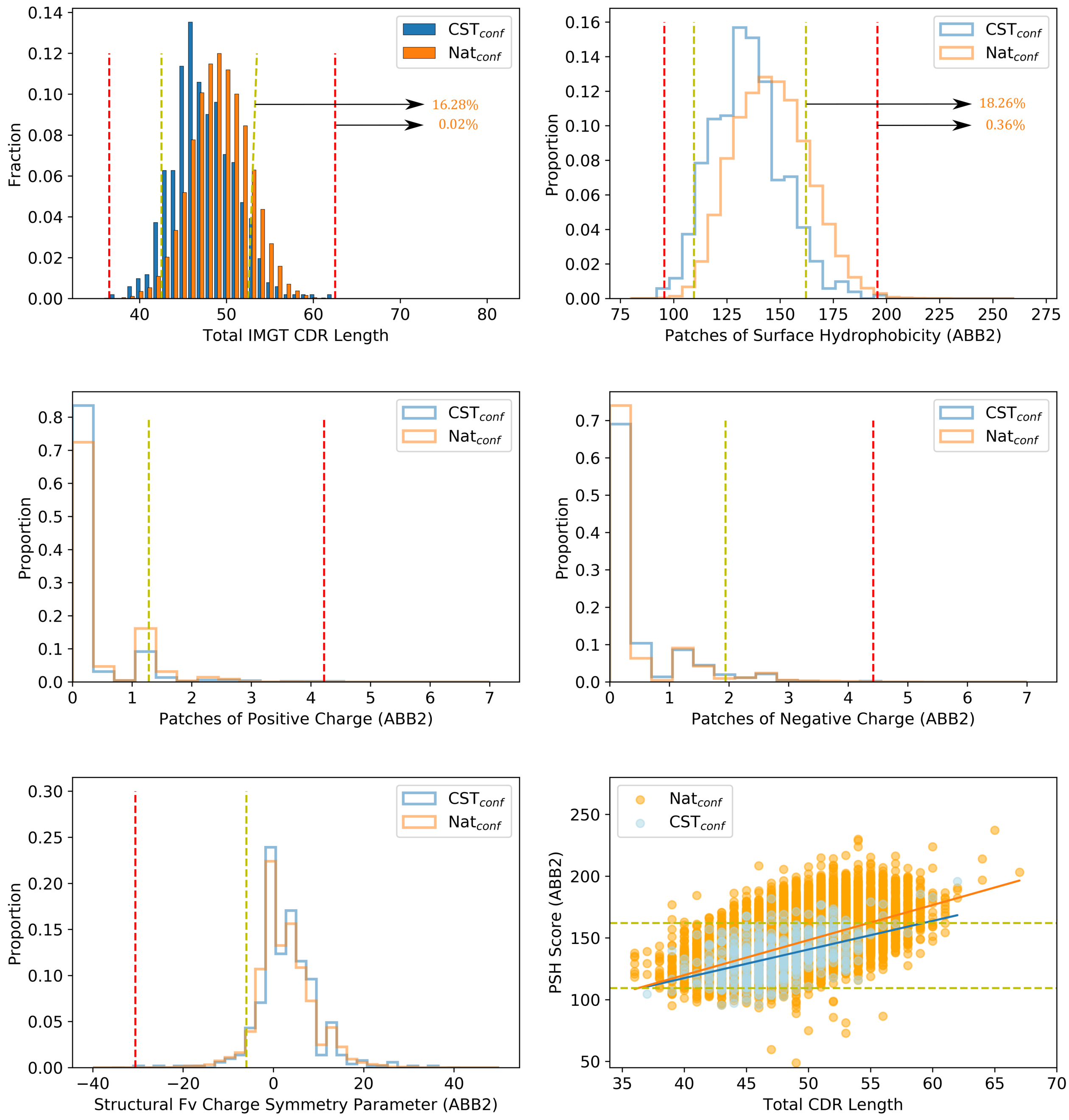


**Fig. 1.** The five TAP developability metrics calculated over the set of CDRH3 confidence-filtered CSTs ($CST_{conf}$, blue) and natural human antibodies ($Nat_{conf}$, orange). Amber and red flagging thresholds are calculated based on the $CST_{conf}$ subset. The percentages of $Nat_{conf}$ antibodies surpassing the upper Total CDR Length and Patches of Surface Hydrophobicity thresholds are highlighted.

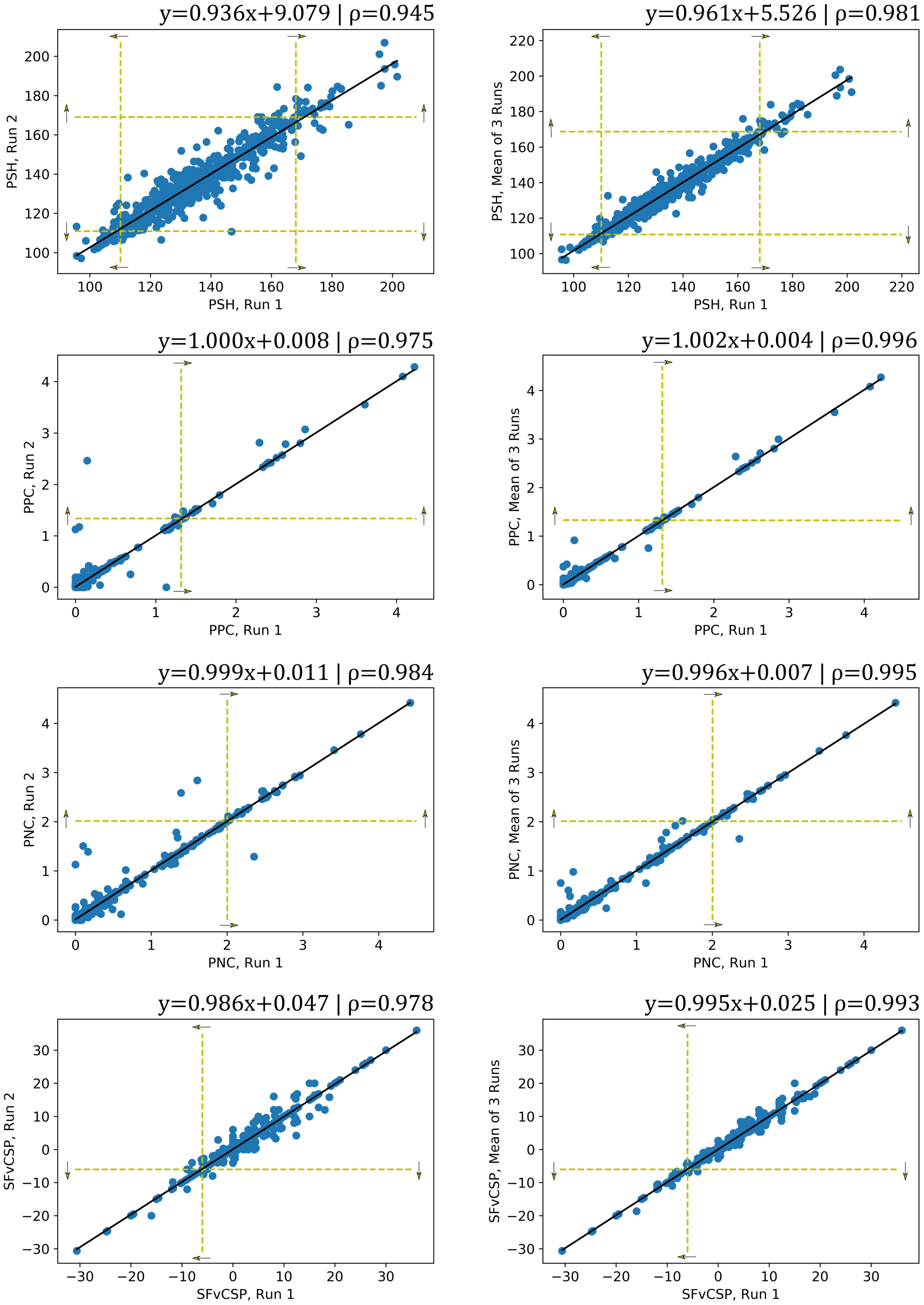


**Fig. 2.** The variation in TAP scores per CST when comparing (left column) one independent ABodyBuilder2 prediction to another, and (right column) one independent ABodyBuilder2 prediction to the mean of three predictions. Amber thresholds are calculated based on the $5^{th}$ and/or $95^{th}$ percentile values for each run/set of runs. Arrows indicate the flagging region relative to each threshold. Least-squares regression lines are plotted with the corresponding equation displayed above each figure.

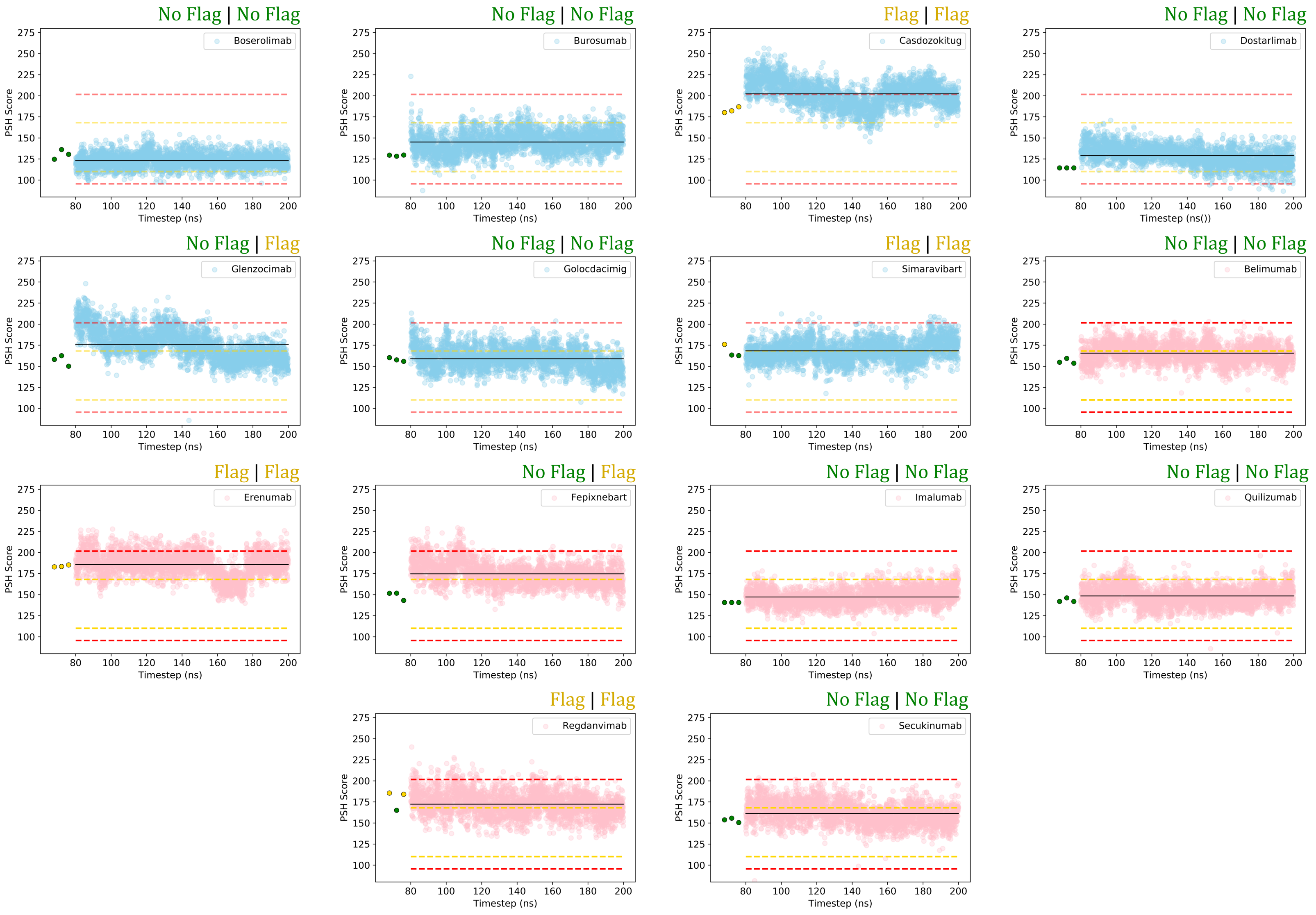


**Fig. 3.** Analysing the Patches of Surface Hydrophobicity (PSH) score for 14 CSTs calculated every 0.04ns over the final 120ns of a 200ns molecular dynamics simulation. The blue trajectories show CSTs that did not have solved Fv structures in the ABodyBuilder2 training set, while pink trajectories show CSTs that did. The amber and red flagging thresholds from Table 1 are shown as dashed lines in corresponding colours; the mean PSH value across the simulation is represented by a solid black line. The PSH values obtained by running TAP on three independent ABodyBuilder2 predictions are shown as spots before the simulation, coloured by the assigned flag. The annotations at the top-right of each graph are in the format [Flag assigned based on the ensemble of three TAP calculations] | [Flag assigned based on the simulation mean value].

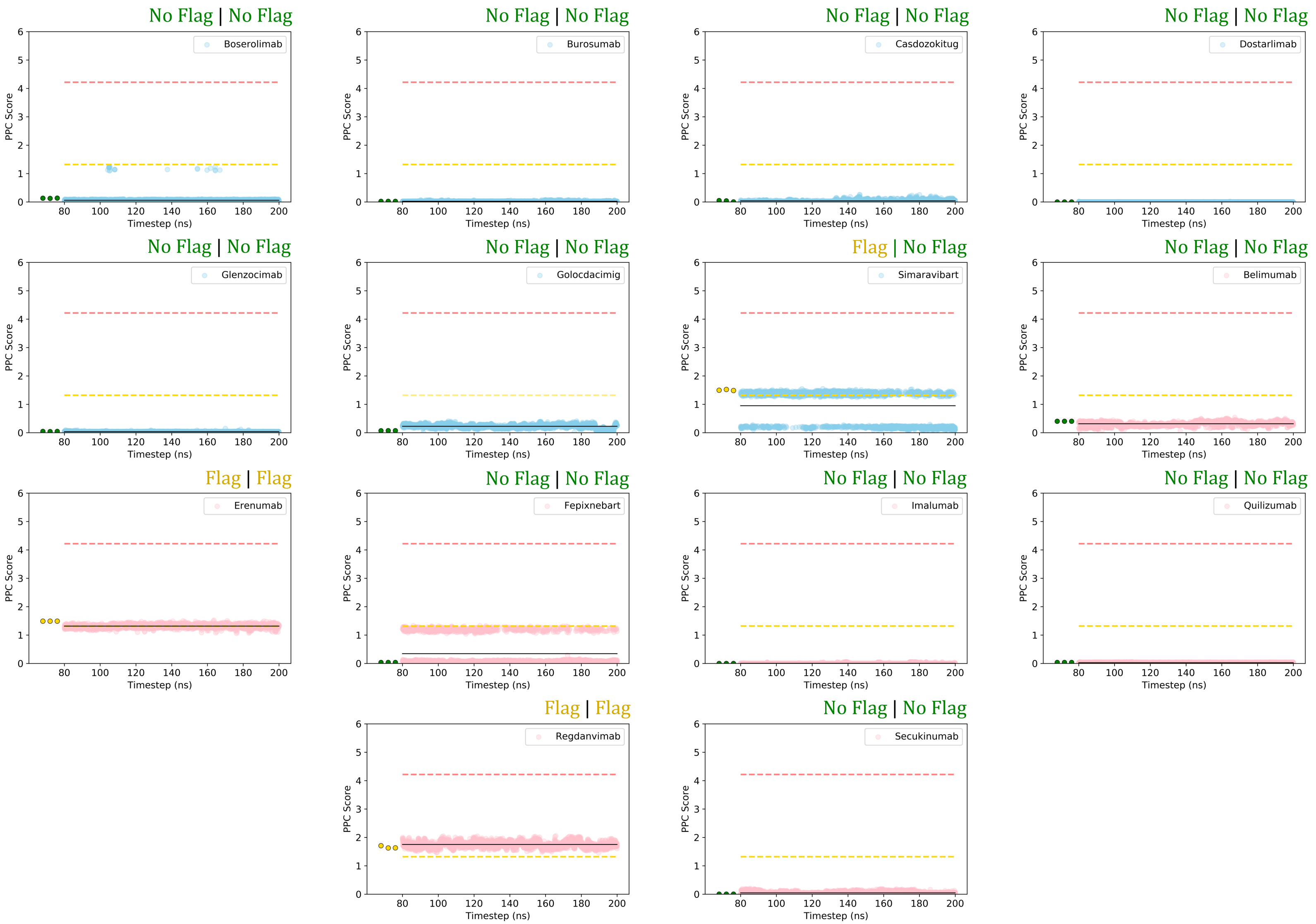


**Fig. 4.** Analysing the Patches of Positive Charge (PPC) score for 14 CSTs calculated every 0.04ns over the final 120ns of a 200ns molecular dynamics simulation. The blue trajectories show CSTs that did not have solved Fv structures in the ABodyBuilder2 training set, while pink trajectories show CSTs that did. The amber and red flagging thresholds from Table 1 are shown as dashed lines in corresponding colours; the mean PSH value across the simulation is represented by a solid black line. The PSH values obtained by running TAP on three independent ABodyBuilder2 predictions are shown as spots before the simulation, coloured by the assigned flag. The annotations at the top-right of each graph are in the format [Flag assigned based on the ensemble of three TAP calculations] | [Flag assigned based on the simulation mean value].

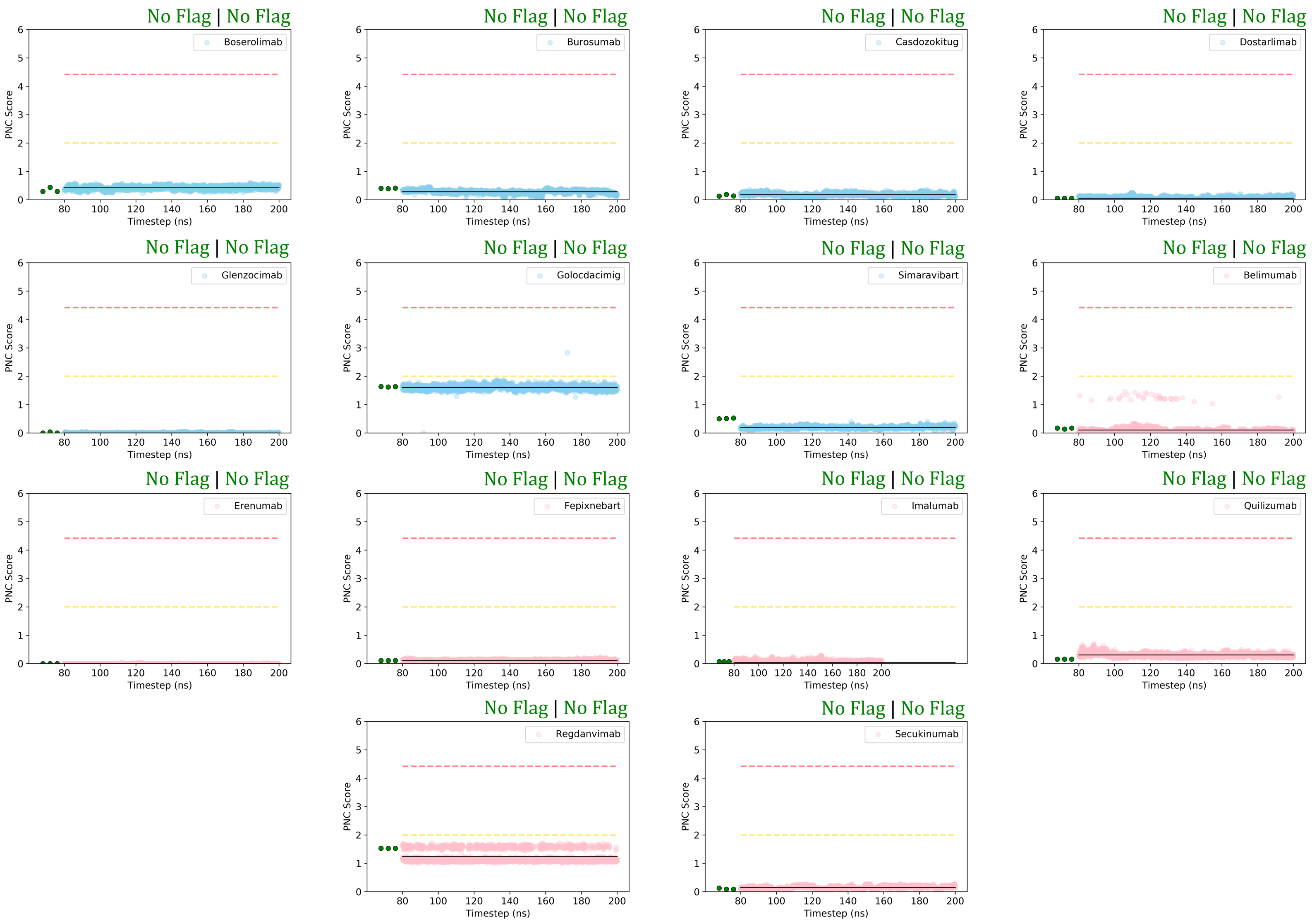


**Fig. 5.** Analysing the Patches of Negative Charge (PNC) score for 14 CSTs calculated every 0.04ns over the final 120ns of a 200ns molecular dynamics simulation. The blue trajectories show CSTs that did not have solved Fv structures in the ABodyBuilder2 training set, while pink trajectories show CSTs that did. The amber and red flagging thresholds from Table 1 are shown as dashed lines in corresponding colours; the mean PSH value across the simulation is represented by a solid black line. The PSH values obtained by running TAP on three independent ABodyBuilder2 predictions are shown as spots before the simulation, coloured by the assigned flag. The annotations at the top-right of each graph are in the format [Flag assigned based on the ensemble of three TAP calculations] | [Flag assigned based on the simulation mean value].

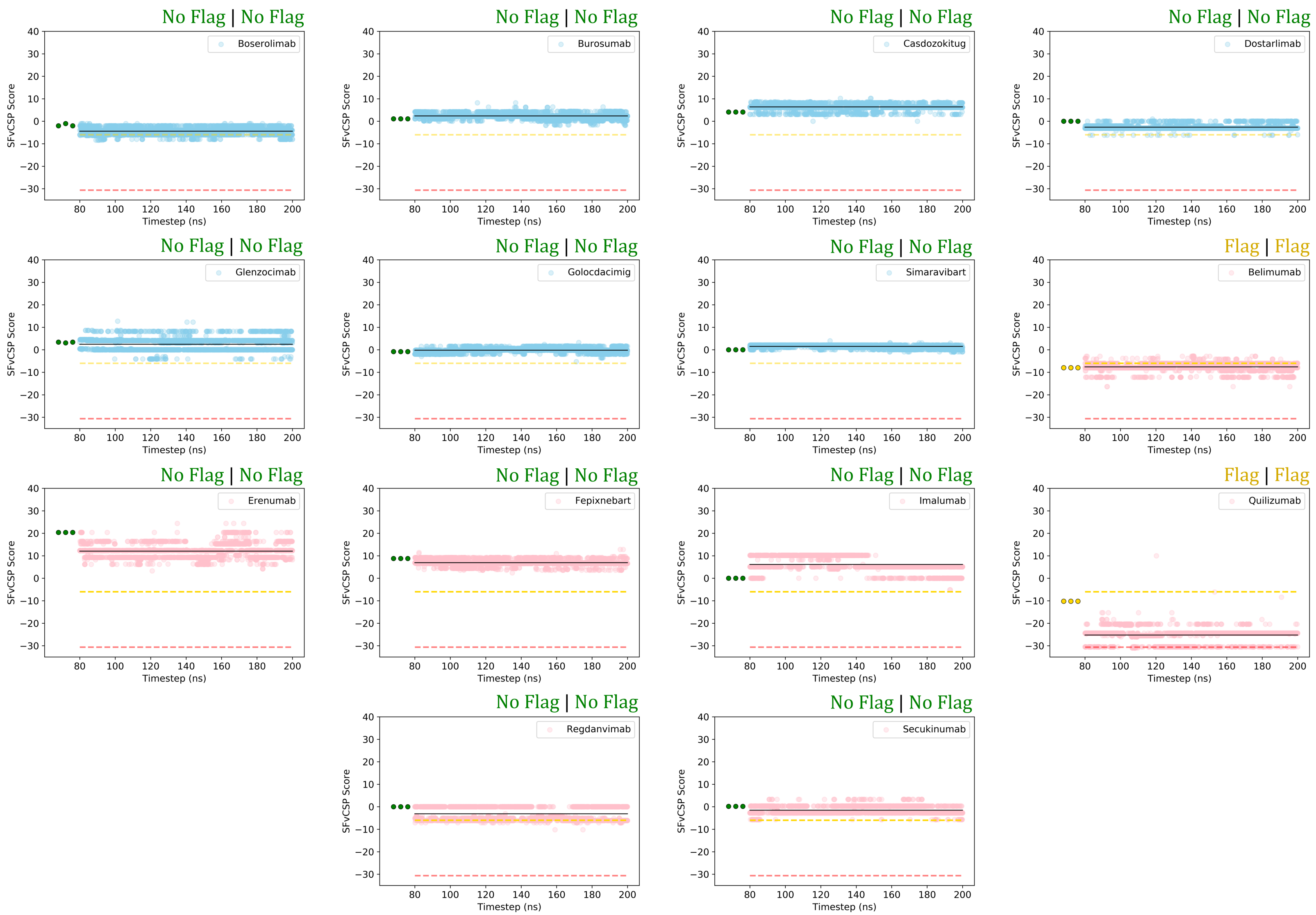


**Fig. 6.** CSTs calculated every 0.04ns over the final 120ns of a 200ns molecular dynamics simulation. The blue trajectories show CSTs that did not have solved Fv structures in the ABodyBuilder2 training set, while pink trajectories show CSTs that did. The amber and red flagging thresholds from Table 1 are shown as dashed lines in corresponding colours; the mean PSH value across the simulation is represented by a solid black line. The PSH values obtained by running TAP on three independent ABodyBuilder2 predictions are shown as spots before the simulation, coloured by the assigned flag. The annotations at the top-right of each graph are in the format [Flag assigned based on the ensemble of three TAP calculations] | [Flag assigned based on the simulation mean value].

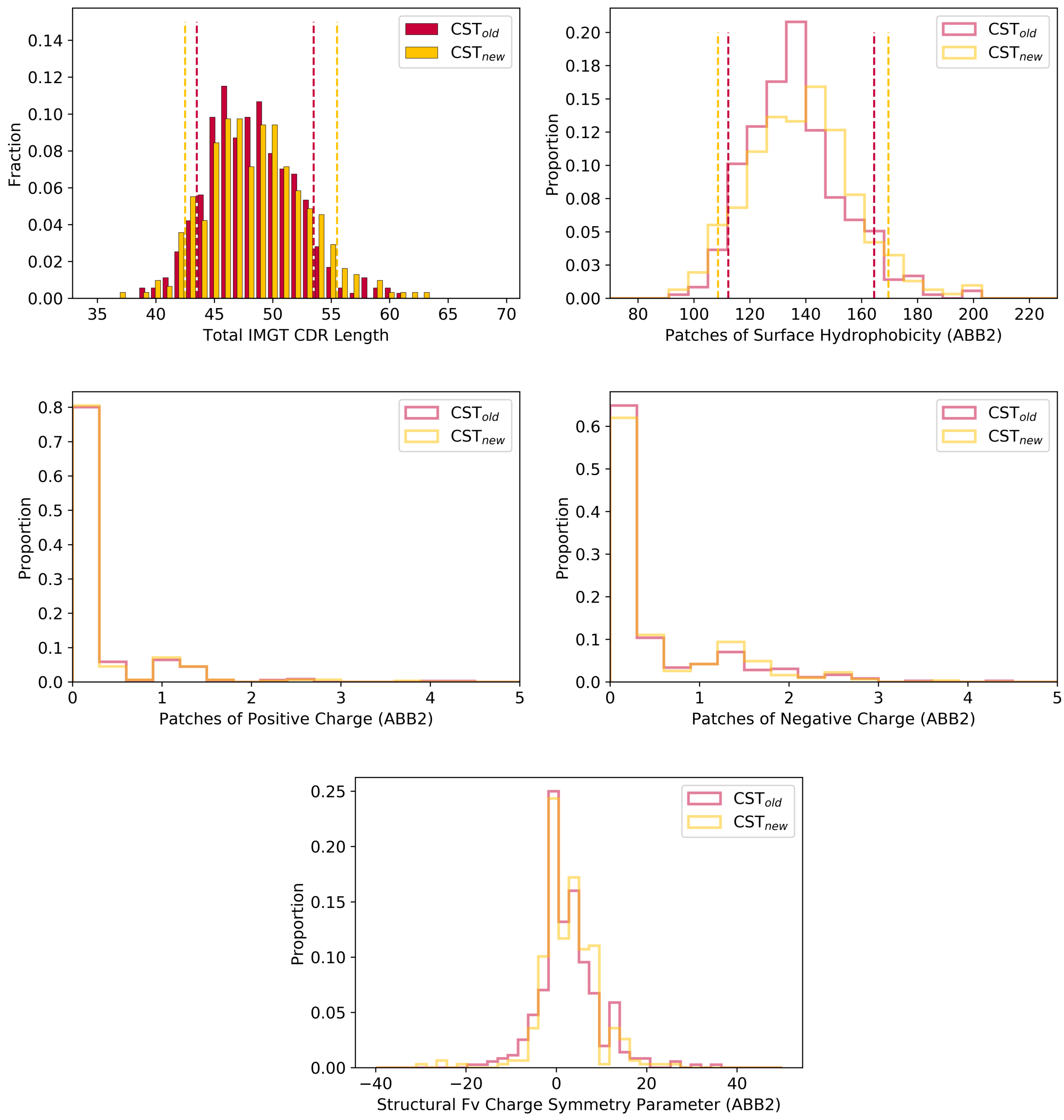


**Fig. 7.** The five TAP developability metrics calculated over the set of CSTs recognised by the WHO between 1987-2017 ($CST_{old}$), and the set recognised between 2018-Present ($CST_{new}$). The amber thresholds for the Total CDR Length and Patches of Surface Hydrophobicity (PSH) properties of each set are highlighted with dashed lines.

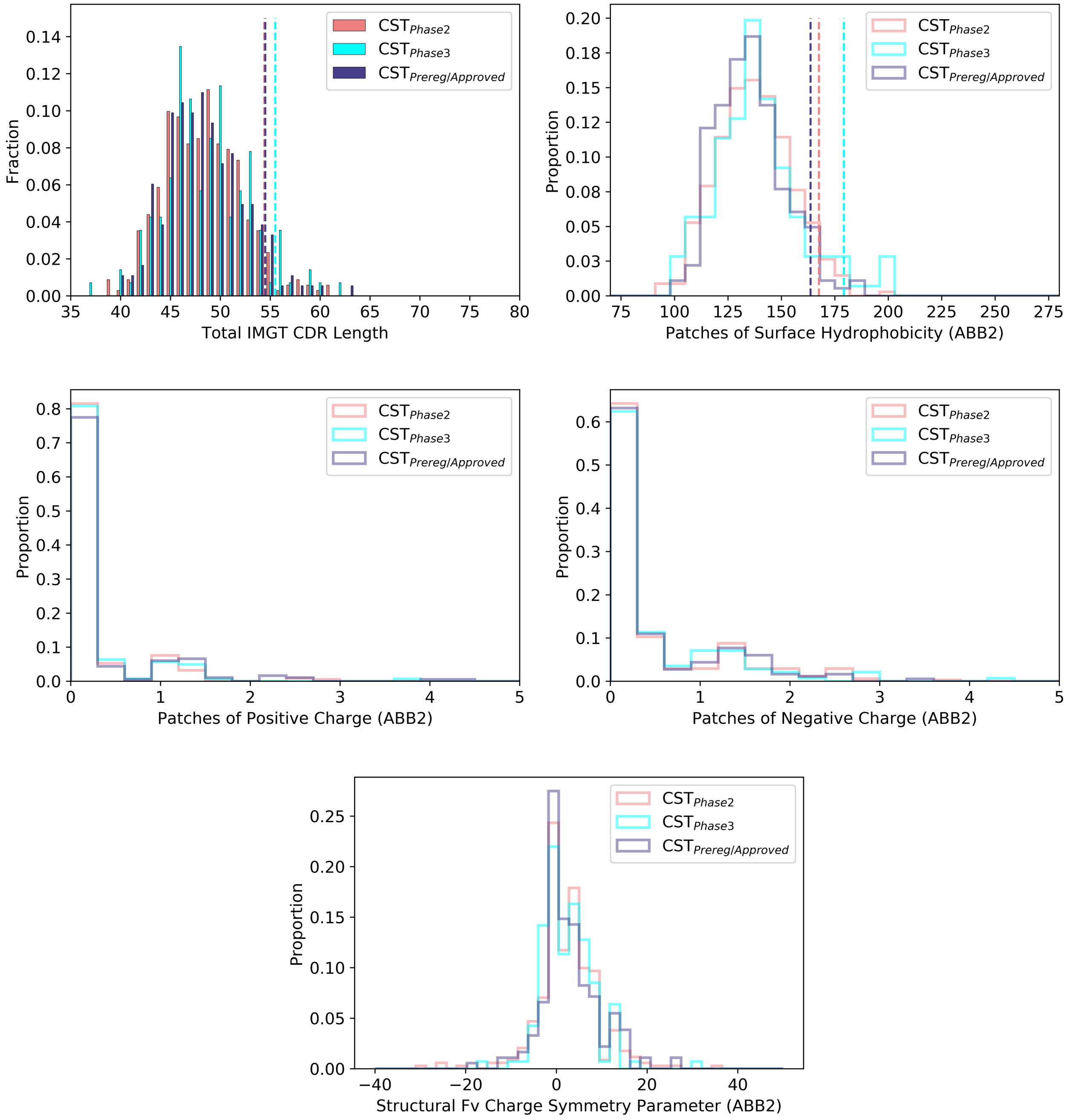


**Fig. 8.** The five TAP developability metrics calculated over the set of CSTs in Phase-II clinical trials (CST$_{\text{Phase2}}$), the set in Phase-III clinical trials (CST$_{\text{Phase3}}$, and the set that have reached Preregistration/been approved as drugs (CST$_{\text{Prereg/Approved}}$). The amber thresholds for the Total CDR Length and Patches of Surface Hydrophobicity (PSH) properties of each set are highlighted with dashed lines.

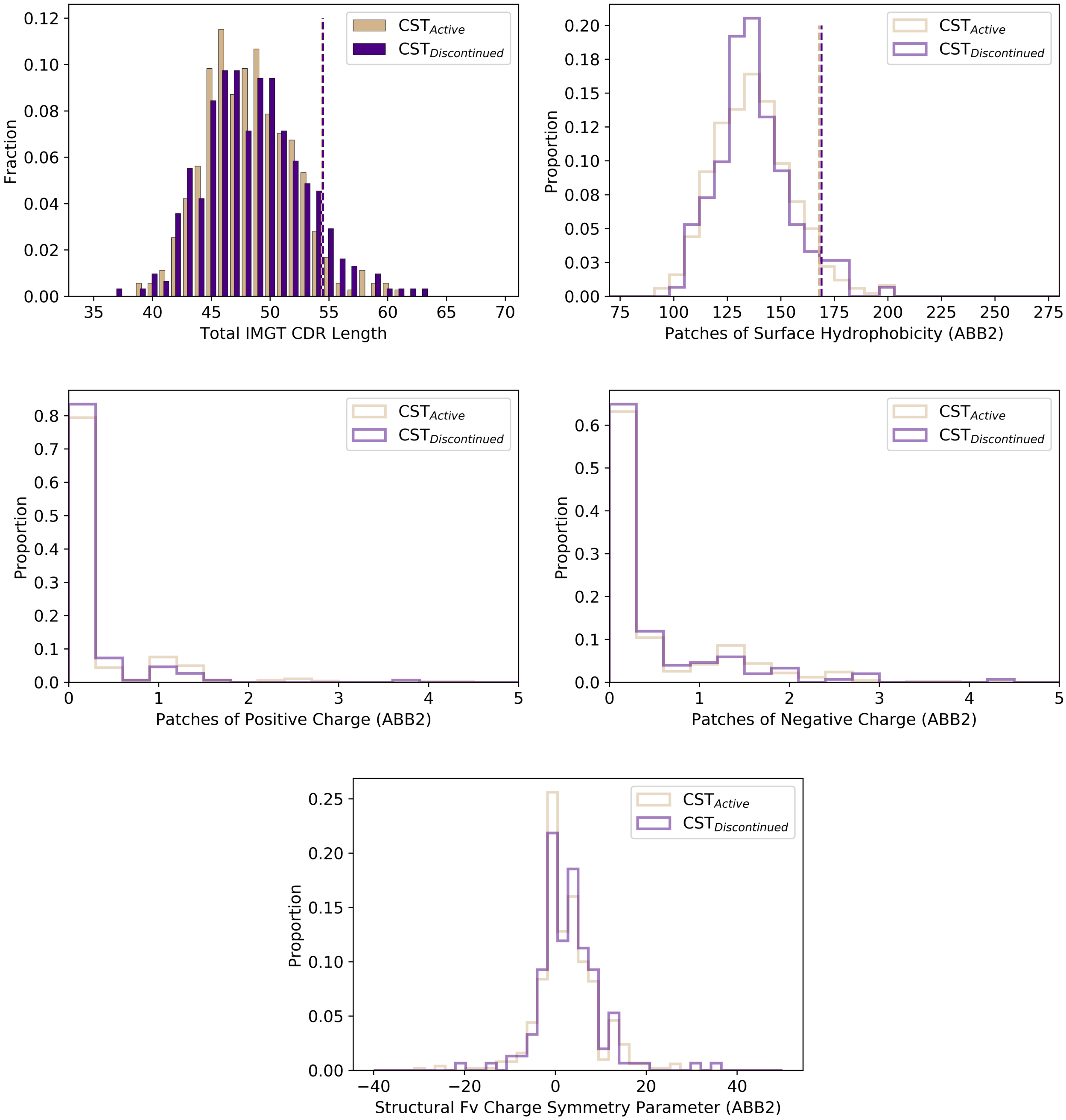


**Fig. 9.** The five TAP developability metrics calculated over the set of CSTs in active development/that completed the development pipeline ($CST_{Active}$) and the set those development campaigns were terminated before approval ($CST_{Discontinued}$. The amber thresholds for the Total CDR Length and Patches of Surface Hydrophobicity (PSH) properties of each set are highlighted with dashed lines.

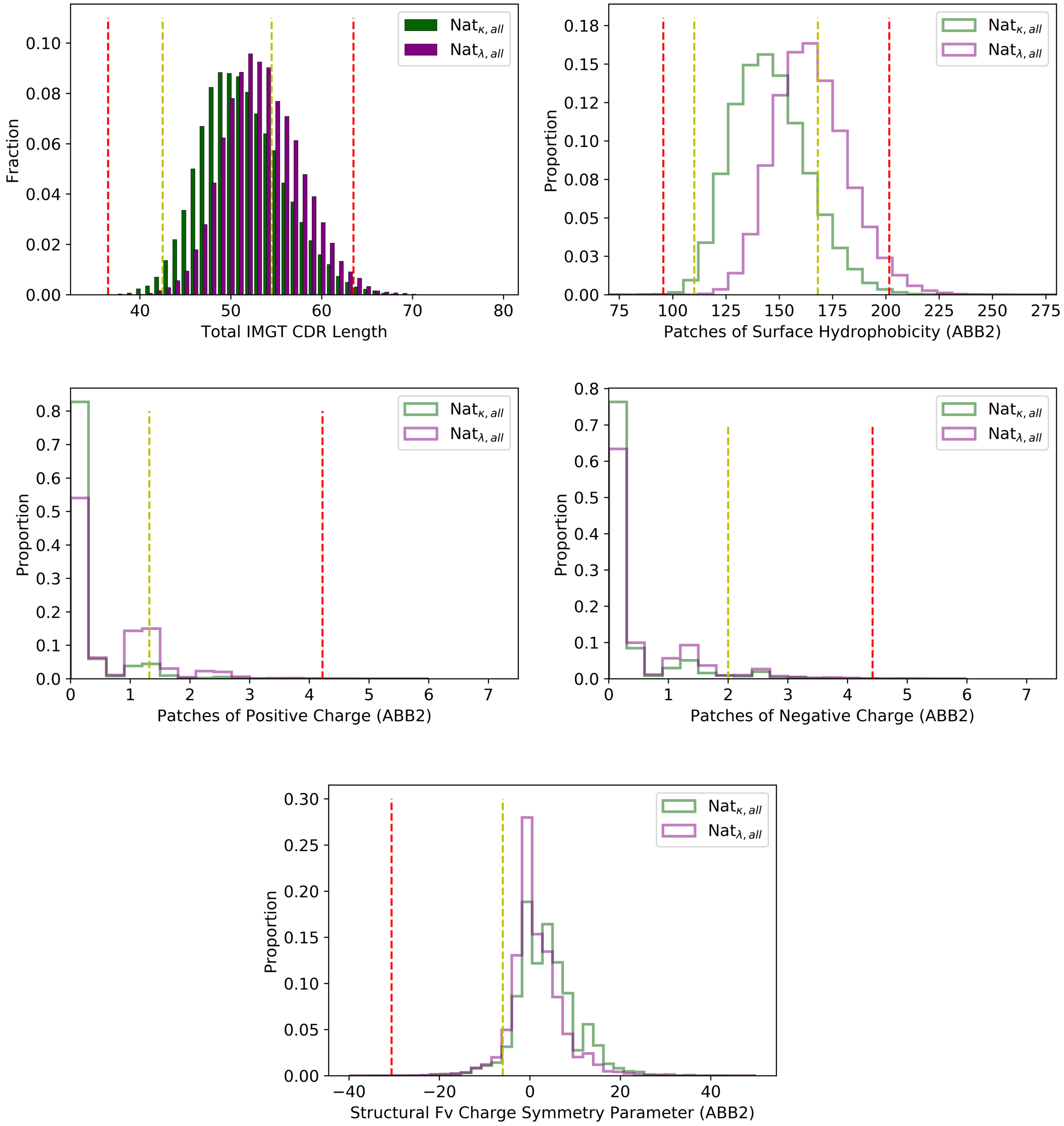


**Fig. 10.** The five TAP developability metrics calculated over all natural human $\kappa$-antibodies and $\lambda$-antibodies.

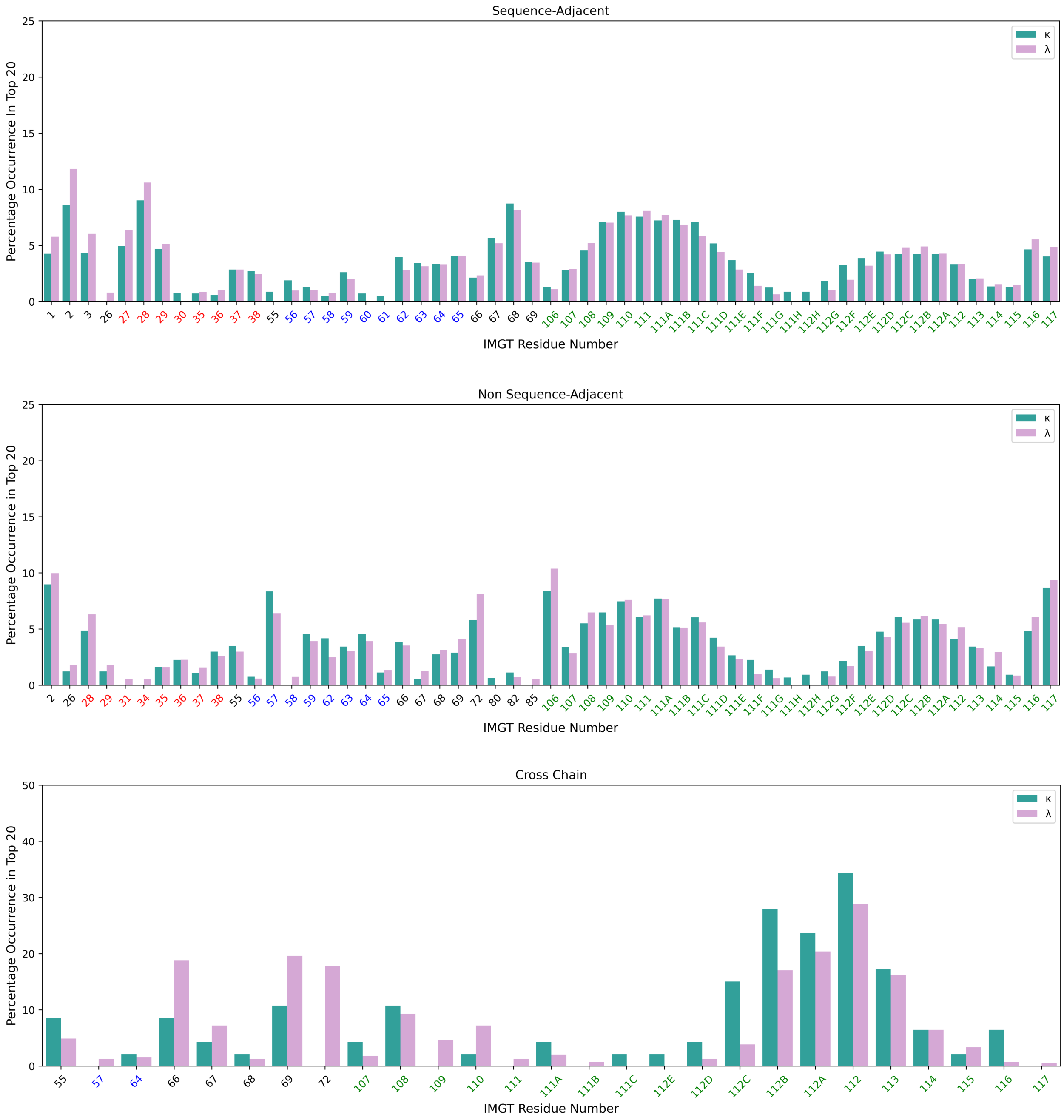


**Fig. 11.** The percentage frequency that each heavy chain IMGT residue position occurs in the top-20-most hydrophobic (above) sequence adjacent and (middle) sequence non-adjacent interactions amongst $\kappa$ (seagreen) and $\lambda$ (plum) red-flagging antibodies. (Bottom) For each heavy chain position, the proportion of top-20-most hydrophobic sequence non-adjacent interactions involving that position that are cross-chain (i.e. involve a light chain residue). Residue numbers in the IMGT-defined CDR1 region are coloured red, in the IMGT-defined CDR2 regions are coloured blue, and in the IMGT-defined CDR3 regions are coloured green.

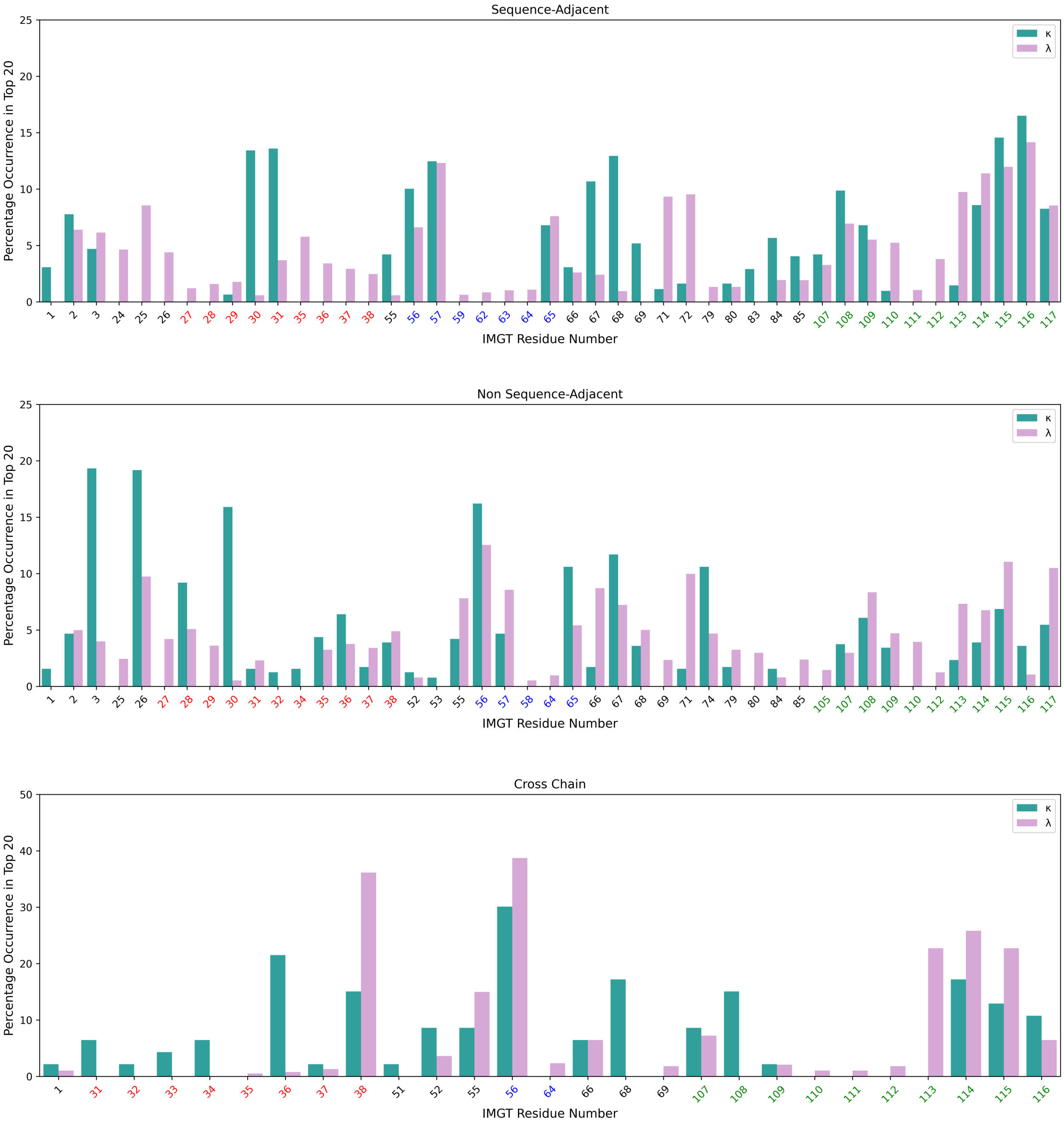


**Fig. 12.** The percentage frequency that each light chain IMGT residue position occurs in the top-20-most hydrophobic (above) sequence adjacent and (middle) sequence non-adjacent interactions amongst $\kappa$ (seagreen) and $\lambda$ (plum) red-flagging antibodies. (Bottom) For each light chain position, the proportion of top-20-most hydrophobic sequence non-adjacent interactions involving that position that are cross-chain (i.e. involve a heavy chain residue). Residue numbers in the IMGT-defined CDR1 region are coloured red, in the IMGT-defined CDR2 regions are coloured blue, and in the IMGT-defined CDR3 regions are coloured green.

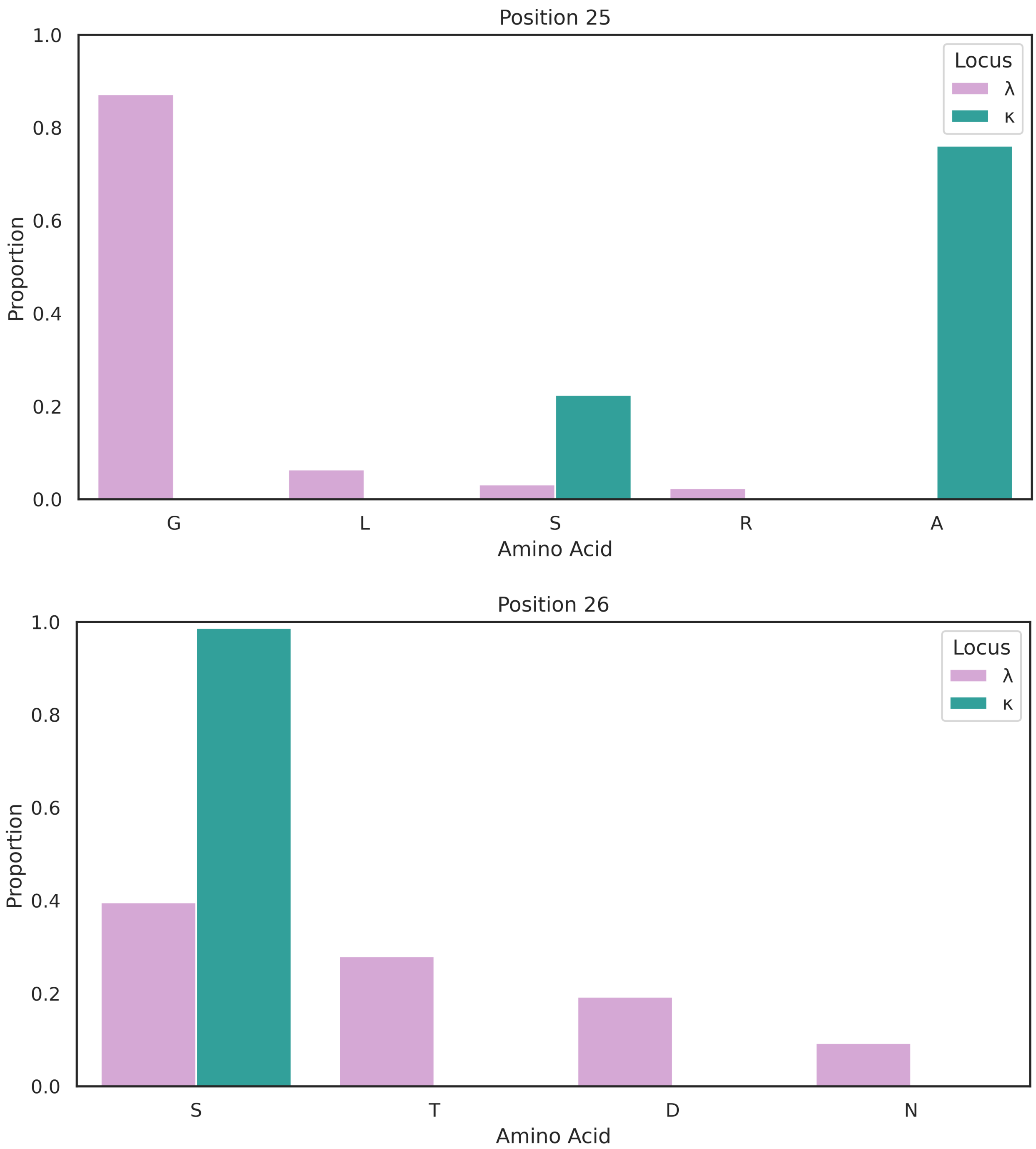


**Fig. 13.** Bar charts showing the amino acid usages at (above) IMGT position 25 and (below) IMGT position 26 amongst natural $\lambda$-antibodies and natural $\kappa$-antibodies.

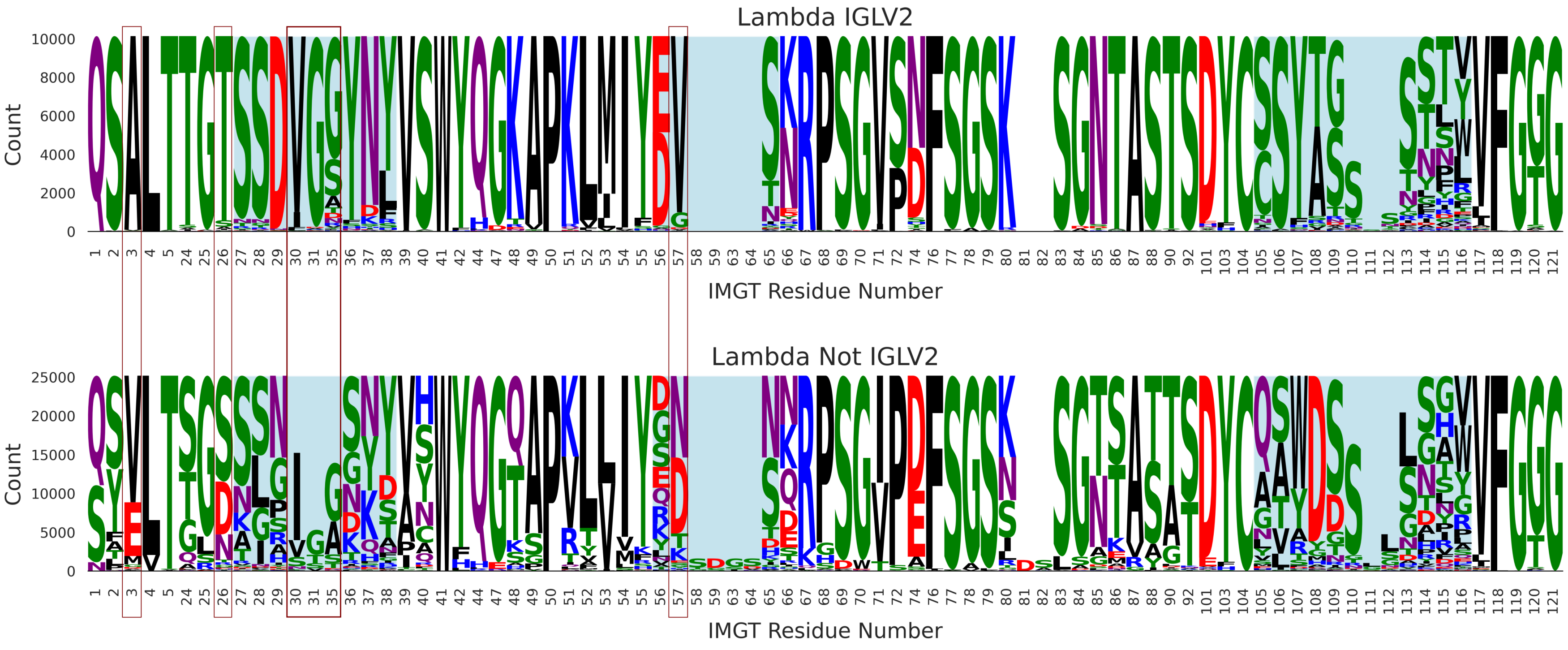


**Fig. 14.** Sequence logo plots showing residue abundance across (top) IGLV2 $\lambda$-antibodies and (bottom) non-IGLV2 $\lambda$-antibodies by IMGT (1) residue position. Red-boxed positions are highlighted in the main manuscript.

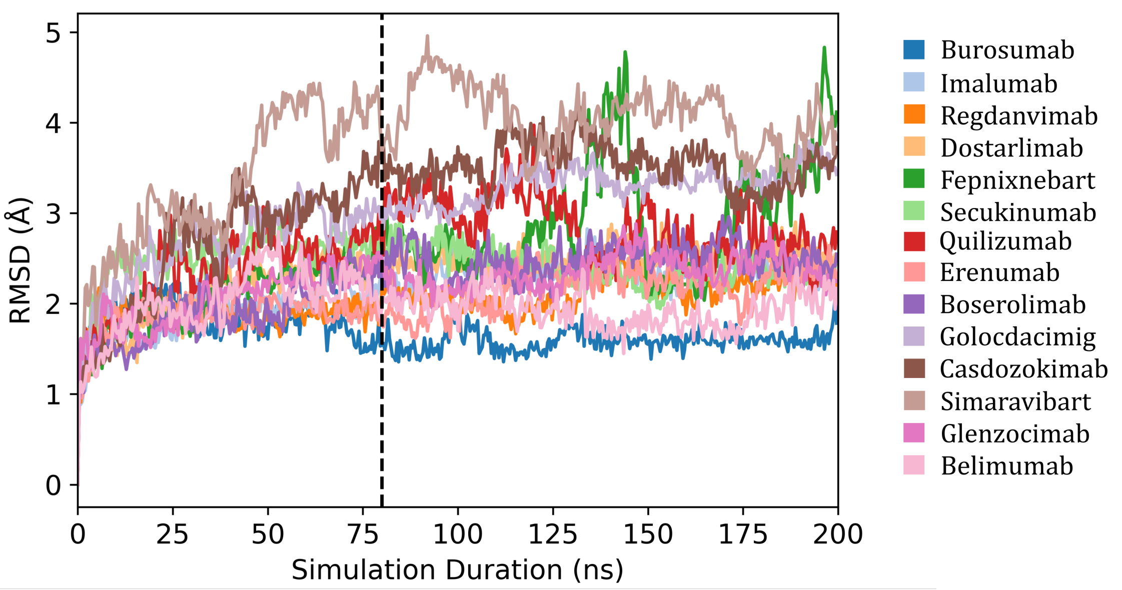


**Fig. 15.** The root-mean squared deviation (RMSD) from the starting structure over the course of the 200 ns simulation for the 14 CSTs studied with molecular dynamics. TAP physicochemical properties were calculated on snapshots post-80 ns, once most simulations had begun to oscillate around a mean RMSD value.

| **Therapeutic** | **PDB ID (Chains)** | **State, $\kappa/\lambda$, Resolution (Å)** |
|---|---|---|
| Boserolimab | 8DS5 (CB) | Complex, $\kappa$, 1.93 |
| Burosumab | 7VEN (BA) | Apo, $\kappa$, 1.45 |
| Casdozokitug | 7ZXK (HL) | Complex, $\kappa$, 2.20 |
| Cemiplimab | 7WVM (AB) | Complex, $\kappa$, 3.40 |
| Dostarlimab | 7WSL (HL) | Complex, $\kappa$, 1.75 |
| Glenzocimab | 7R58 (HL) | Complex, $\kappa$, 1.90 |
| Golocdacimig | 7R8U (HL) | Complex, $\lambda$, 1.90 |
| Simaravibart | 7SBU (HL) | Complex, $\kappa$, 2.53 |

| | Mean RMSD (Å) | | | | | | | |
|---|---|---|---|---|---|---|---|---|
| | H1 | H2 | H3 | L1 | L2 | L3 | % All S.E.R. Correct | % CDR S.E.R. Correct |
| ABodyBuilder1 | 1.73 | 1.00 | 3.32 | 0.91 | 0.56 | 1.16 | 95.99% | 94.14% |
| ABodyBuilder2 | 0.78 | 0.95 | 2.68 | 0.77 | 0.50 | 0.80 | 96.39% | 94.87% |
| % Improvement | *54.9%* | *5.0%* | *19.3%* | *5.4%* | *10.7%* | *31.0%* | *+0.40%* | *+0.73%* |

**Table 1.** (Above) The eight clinical-stage therapeutics (CSTs) with 100% sequence identity solved crystal structures publicly released after 31$^{st}$ July 2021 (ABodyBuilder2's training set cutoff date). (Below) The performance of ABodyBuilder1 (5) *vs.* ABodyBuilder2 (2) on this subset of CSTs. CDR: Complementarity-determining region; S.E.R: Surface exposed residues.

| | | | |
|---|---|---|---|
| Abituzumab ($\lambda$) | Adalimumab ($\kappa$) | Aducanumab ($\kappa$) | Alemtuzumab ($\kappa$) |
| Alomfilimab ($\kappa$) | Amivantamab ($\kappa$) | Amubarvimab ($\kappa$) | Andecalizumab ($\kappa$) |
| Anifrolumab ($\kappa$) | Arcitumumab ($\kappa$) | Atezolizumab ($\kappa$) | Avelumab ($\lambda$) |
| Bamlanivimab ($\kappa$) | Basiliximab ($\kappa$) | Bebtelovimab ($\lambda$) | Belimumab ($\lambda$) |
| Bentracimab ($\lambda$) | Benufutamab ($\kappa$) | Berlimatoxumab ($\kappa$) | Bevacizumab ($\kappa$) |
| Bezlotoxumab ($\kappa$) | Bimagrumab ($\lambda$) | Bococizumab ($\kappa$) | Briakinumab ($\lambda$) |
| Camrelizumab ($\kappa$) | Certolizumab ($\kappa$) | Cetuximab ($\kappa$) | Cinpanemab ($\lambda$) |
| Clesrovimab ($\kappa$) | Coltuximab ($\kappa$) | Conatumumab ($\kappa$) | Concizumab ($\kappa$) |
| Crenezumab ($\kappa$) | Crovalimab ($\kappa$) | Daclizumab ($\kappa$) | Daratumumab ($\kappa$) |
| Diridavumab ($\lambda$) | Dupilumab ($\kappa$) | Durvalumab ($\kappa$) | Eculizumab ($\kappa$) |
| Efalizumab ($\kappa$) | Emactuzumab ($\kappa$) | Erenumab ($\lambda$) | Erlizumab ($\kappa$) |
| Etesevimab ($\kappa$) | Fepixnebart ($\kappa$) | Gantenerumab ($\kappa$) | Gevokizumab ($\kappa$) |
| Golimumab ($\kappa$) | Guselkumab ($\lambda$) | Ibalizumab ($\kappa$) | Ibritumumomab ($\kappa$) |
| Idarucizumab ($\kappa$) | Imalumab ($\kappa$) | Infliximab ($\kappa$) | Ipilimumab ($\kappa$) |
| Isatuximab ($\kappa$) | Ixekizumab ($\kappa$) | Izalontamab ($\kappa$) | Lampalizumab ($\kappa$) |
| Lanadelumab ($\kappa$) | Lebrikizumab ($\kappa$) | Ligelizumab ($\kappa$) | Lumretuzumab ($\kappa$) |
| Matuzumab ($\kappa$) | Metelimumab ($\kappa$) | Mevonlerbart ($\kappa$) | Motavizumab ($\kappa$) |
| Muromonab ($\kappa$) | Necitumumab ($\kappa$) | Nirsevimab ($\kappa$) | Nivolumab ($\kappa$) |
| Obinutuzumab ($\kappa$) | Ofatumumab ($\kappa$) | Ogalvibart ($\kappa$) | Olokizumab ($\kappa$) |
| Omalizumab ($\kappa$) | Omburtamab ($\kappa$) | Ontamalimab ($\kappa$) | Opicinumab ($\kappa$) |
| Orilanolimab ($\kappa$) | Panitumumab ($\kappa$) | Paridiprubart ($\kappa$) | Pateclizumab ($\kappa$) |
| Pembrolizumab ($\kappa$) | Pertuzumab ($\kappa$) | Ponezumab ($\kappa$) | Prezalumab ($\kappa$) |
| Quilizumab ($\kappa$) | Radretumab ($\kappa$) | Ramucirumab ($\kappa$) | Ranibizumab ($\kappa$) |
| Ravagalimab ($\kappa$) | Regdanvimab ($\lambda$) | Rituximab ($\kappa$) | Rontalizumab ($\kappa$) |
| Rozanolixizumab ($\kappa$) | Ruplizumab ($\kappa$) | Secukinumab ($\kappa$) | Serplulimab ($\kappa$) |
| Sifalimumab ($\kappa$) | Spesolimab ($\kappa$) | Suvratoxumab ($\kappa$) | Talacotuzumab ($\kappa$) |
| Tanezumab ($\kappa$) | Teneliximab ($\kappa$) | Tezepelumab ($\lambda$) | Tislelizumab ($\kappa$) |
| Tixagevimab ($\kappa$) | Toripalimab ($\kappa$) | Tralokinumab ($\lambda$) | Trastuzumab ($\kappa$) |
| Tremlimumab ($\kappa$) | Urelumab ($\kappa$) | Ustekinumab ($\kappa$) | Utomilumab ($\lambda$) |
| Vanucizumab ($\lambda$) | Vonlerolizumab ($\kappa$) | Zenocutuzumab ($\kappa$) | |

**Table 2.** The 119 clinical-stage therapeutics (CSTs, 103 x $\kappa$, 16 x $\lambda$) with 100% sequence identity solved crystal structures publicly released on or before 31$^{st}$ July 2021 (ABodyBuilder2's training set cutoff date). Models of these therapeutics were excluded in benchmarking studies. Res: Resolution.

| TAP Property | Amber Flag Region | Red Flag Region |
|---|---|---|
| Total CDR Length ($L_{tot}$) | 37 (*0*) $\leq L_{tot} \leq$ 42 (*0*)<br>53 (-*2*) $\leq L_{tot} \leq$ 62 (-*1*) | $L_{tot} <$ 37 (*0*)<br>$L_{tot} >$ 62 (-*1*) |
| Patches of Surface Hydrophobicity (PSH) | 95.58 (*0*) $\leq$ PSH $\leq$ 109.51 (-*0.60*)<br>162.15 (-*5.91*) $\leq$ PSH $\leq$ 195.68 (-*5.91*) | PSH $<$ 95.58 (*0*)<br>PSH $>$ 195.68 (-*5.91*) |
| Patches of Positive Charge (PPC) | 1.28 (-*0.04*) $\leq$ PPC $\leq$ 4.22 (-*0.13*) | PPC $>$ 4.22 (-*0.13*) |
| Patches of Negative Charge (PNC) | 1.94 (-*0.06*) $\leq$ PNC $\leq$ 4.42 (*0*) | PNC $>$ 4.42 (*0*) |
| Structural Fv Charge Symmetry Parameter (SFvCSP) | -30.6 (*0*) $\leq$ SFvCSP $\leq$ -6.0 (*0*) | SFvCSP $<$ -30.6 (*0*) |

**Table 3.** Flagging regions across the five TAP developability metrics calculated over the 510 CSTs that are modeled with higher CDRH3 confidence (*i.e.* the $CST_{conf}$ dataset). Differences from the $CST_{all}$ guidelines are provided in the brackets.

| Metric | All, Mean Variance/3 Runs | $\kappa$, Mean Variance/3 Runs | $\lambda$, Mean Variance/3 Runs |
|---|---|---|---|
| PSH | 10.533 | 10.455 | 11.046 |
| PPC | 0.004 | 0.004 | 0.000 |
| PNC | 0.005 | 0.005 | 0.007 |
| SFvCSP | 0.572 | 0.597 | 0.404 |

**Table 4.** The mean variance recorded for each structure-based TAP developability metric calculated on three repeat ABodyBuilder2 models of all 664 CSTs (column 2), the subset of 576 $\kappa$-CSTs only (column 3), and the subset of 88 $\lambda$-CSTs only (column 4).

| TAP Property | Amber Flag Region | Red Flag Region |
|---|---|---|
| Patches of Surface Hydrophobicity (PSH) | 94.85 (-*0.73*) $\leq$ PSH $\leq$ 110.78 (+*0.67*)<br>168.74 (+*0.68*) $\leq$ PSH $\leq$ 206.93 (+*5.34*) | PSH $<$ 94.85 (-*0.73*)<br>PSH $>$ 206.93 (+*5.34*) |
| Patches of Positive Charge (PPC) | 1.33 (+*0.01*) $\leq$ PPC $\leq$ 4.31 (+*0.09*) | PPC $>$ 4.31 (+*0.09*) |
| Patches of Negative Charge (PNC) | 2.01 (+*0.01*) $\leq$ PNC $\leq$ 4.42 (*0*) | PNC $>$ 4.42 (*0*) |
| Structural Fv Charge Symmetry Parameter (SFvCSP) | -30.60 (*0*) $\leq$ SFvCSP $\leq$ -6.00 (*0*) | SFvCSP $<$ -30.60 (*0*) |

**Table 5.** The TAP developability guidelines for structure-dependent metrics set by combining three repeat modeling runs for each of the 664 CST Fvs (2). Differences from the $CST_{all}$ guidelines are provided in the brackets.

| CST | PDB Structure Used for Constant Region Grafting (chain IDs, heavy + light) |
|---|---|
| Belimumab | 5Y9K (HL) |
| Boserolimab | 8DS5 (CB) |
| Burosumab | 7VEN (BA) |
| Casdozokitug | 7ZXK (HL) |
| Dostarlimab | 7WSL (HL) |
| Erenumab | 6UMH (HL) |
| Fepnixnebart | 5KN5 (AB) |
| Glenzocimab | 7R58 (HL) |
| Golocdacimig | 7R8U (HL) |
| Imalumab | 6FOE (HL) |
| Quilizumab | 3HR5 (HL) |
| Regdanvimab | 7CM4 (HL) |
| Secukinumab | 6WIO (AB) |
| Simaravibart | 7SBU (HL) |

**Table 6.** Protein Data Bank IDs and chain IDs mapping to the coordinates used for CH1/CL domain grafting for each clinical-stage therapeutic (CST).

| Stage | Ensemble | Restrained Atoms | Restraint Strength (kJ mol$^{-1}$ nm$^{-2}$) | Duration | Start T (K) | End T (K) |
|---|---|---|---|---|---|---|
| 1 | Minimisation | Protein Heavy | 4184.00 | 5000 steps | - | - |
| 2 | NVT | Protein Heavy | 4184.00 | 0.2 ns | 100 | 300 |
| 3 | NPT | Protein Heavy | 4184.00 | 0.2ns | 300 | 300 |
| 4 | NPT | Protein Heavy | 2092.00 | 0.5ns | 300 | 300 |
| 5 | Minimisation | Backbone Heavy | 2092.00 | 5000 steps | 300 | 300 |
| 6 | NPT | Backbone Heavy | 2092.00 | 0.2ns | 300 | 300 |
| 7 | NPT | Backbone Heavy | 418.40 | 0.2ns | 300 | 300 |
| 8 | NPT | Backbone Heavy | 41.84 | 0.2ns | 300 | 300 |
| 9 | NPT | - | - | 1ns | 300 | 300 |

**Table 7.** Equilibration protocol steps. Where start and end temeperatures differ, the temperature was linearly increased over the duration of the stage. T: Temperature.

| Human Gene Family | # CSTs | % Abundance Amongst these CSTs | % Abundance Amongst Natural Sequences |
|---|---|---|---|
| LV1 | 24 | 33.80 | 29.73 |
| LV2 | 15 | 21.13 | 28.71 |
| LV3 | 25 | 35.21 | 28.11 |
| LV4 | 0 | 0 | 3.08 |
| LV5 | 0 | 0 | 0.94 |
| LV6 | 3 | 4.23 | 3.08 |
| LV7 | 3 | 4.23 | 3.17 |
| LV8 | 1 | 1.41 | 1.82 |
| LV9 | 0 | 0 | 0.55 |
| LV10 | 0 | 0 | 0.80 |

**Table 8.** Comparison of gene family usages across human gene-derived $\lambda$-CSTs, and gene family uses across natural paired sequences from OAS (6).

| Human Gene | # CSTs | % Abundance Amongst these CSTs | % Abundance Amongst LV2 Gene-Derived Natural Sequences |
|---|---|---|---|
| LV2-8 | 0 | 0 | 13.83 |
| LV2-11 | 3 | 20 | 11.62 |
| LV2-14 | 10 | 66.67 | 42.85 |
| LV2-18 | 0 | 0 | 13.83 |
| LV2-23 | 2 | 13.33 | 17.87 |

**Table 9.** Comparison of gene usages across human LV2 gene family-derived $\lambda$-CSTs, and gene uses across human LV2 gene family-derived natural paired sequences from OAS (6).

### Bibliography


1. Marie-Paule Lefranc, Christelle Pommié, Manuel Ruiz, Véronique Giudicelli, Elodie Foulquier, Lisa Truong, Valérie Thouvenin-Contet, and Gérard Lefranc. IMGT unique numbering for immunoglobulin and T cell receptor variable domains and Ig superfamily V-like domains. *Dev Comp Immunol*, 27(1):55–77, 2003. doi: 10.1016/s0145-305x(02)00039-3.
2. Brennan Abanades, Wing Ki Wong, Fergus Boyles, Guy Georges, Alexander Bujotzek, and Charlotte M. Deane. ImmuneBuilder: Deep-Learning models for predicting the structures of immune proteins. *Commun Biol*, 6:575, 2023. doi: 110.1038/s42003-023-04927-7.
3. A. Shrake and J. A. Rupley. Environment and exposure to solvent of protein atoms. Lysozyme and insulin. *J Mol Biol*, 79(2):351–371, 1973. doi: 10.1016/0022-2836(73)90011-9.
4. Matthew I. J. Raybould, Claire Marks, Konrad Krawczyk, Bruck Taddese, Jaroslaw Nowak, Alan P. Lewis, Alexander Bujotzek, Jiye Shi, and Charlotte M. Deane. Five computational developability guidelines for therapeutic antibody profiling. *Proc Natl Acad Sci*, 116(10):4025–4030, 2019. doi: 10.1073/pnas.1810576116.
5. Jinwoo Leem, James Dunbar, Guy Georges, Jiye Shi, and Charlotte M. Deane. ABodyBuilder: Automated antibody structure prediction with data–driven accuracy estimation. *mAbs*, 8(7):1259–1268, 2016. doi: 10.1080/19420862.2016.1205773.
6. Tobias H. Olsen, Fergus Boyles, and Charlotte M. Deane. Observed Antibody Space: A diverse database of cleaned, annotated, and translated unpaired and paired antibody sequences. *Protein Sci*, 31(1):141–146, 2022. doi: 10.1002/pro.4205.
